# EpiGraph: an open-source platform to quantify epithelial organization

**DOI:** 10.1101/217521

**Authors:** Pablo Vicente-Munuera, Pedro Gómez-Gálvez, Robert J. Tetley, Cristina Forja, Antonio Tagua, Marta Letrán, Melda Tozluoglu, Yanlan Mao, Luis M. Escudero

## Abstract

During development, cells must coordinate their differentiation with their growth and organization to form complex multicellular structures such as tissues and organs. Healthy tissues must maintain these structures during homeostasis. Epithelia are packed ensembles of cells from which the different tissues of the organism will originate during embryogenesis. A large barrier to the analysis of the morphogenetic changes in epithelia is the lack of simple tools that enable the quantification of cell arrangements. Here we present EpiGraph, an image analysis tool that quantifies epithelial organization. Our method combines computational geometry and graph theory to measure the degree of order of any packed tissue. EpiGraph goes beyond the traditional polygon distribution analysis, capturing other organizational traits that improve the characterization of epithelia. EpiGraph can objectively compare the rearrangements of epithelial cells during development and homeostasis to quantify how the global ensemble is affected. Importantly, it has been implemented in the open-access platform FIJI. This makes EpiGraph very user friendly, with no programming skills required.

## INTRODUCTION

The development of any multicellular organism is based on coordinated changes that transform the embryo into the adult individual. During morphogenesis and growth, patterning, cell divisions and architectural changes must perfectly fit together for the correct development of the body plan. Any morphogenetic movement such as migration, extension or invagination of epithelial cells is coupled with dramatic changes in the organization of cells (Bertet et al., 2004; Blankenship et al., 2006; Escudero et al., 2007; Farhadifar et al., 2007; Girdler and Roper, 2014; Gómez-Gálvez et al., 2018; Lecuit and Lenne, 2007; Pilot and Lecuit, 2005). After development, homeostatic tissues must maintain their complex organization of cells in order to function correctly.

How tissues modulate and maintain their organization during development and homeostasis is an important question that remains unsolved. This is mainly due to the lack of simple and general methods that can capture and quantify the arrangement of cells. It has been known for almost a hundred years that epithelial tissues exhibit a degree of order. The analysis of epithelial organization has been mainly based on the number of neighbours of the epithelial cells, considering the apical surface of these cells as convex polygons with the same number of sides as neighbours. In previous works, we have investigated several aspects of the organization of packed tissues using Voronoi tessellations to compare the polygon distributions of natural and mathematical tessellations (Sanchez-Gutierrez et al., 2016). We have described that the polygon distribution of natural tessellations is restricted to a series of frequencies of polygons that match the Voronoi diagrams that conform to the Centroidal Voronoi tessellation (CVT). This is what we call a “CVT path” and was used as a scale to compare the organization of different packed tissues. However, polygon distribution is not sufficient to completely characterize tissue organization. Tissues with clearly different appearance can present very similar polygon distribution (Sanchez-Gutierrez et al., 2016).

As an alternative approach, we have proposed that Graph Theory could capture differences in the topology of tissues (Escudero et al., 2011; Sanchez-Gutierrez et al., 2013; Sánchez-Gutiérrez et al., 2017). This is based on the idea of converting the epithelium into a network of cell-to-cell contacts (Escudero et al., 2011). The resulting “epithelial graph” can be analysed by combining the tools of network theory and multivariable statistical analysis (Escudero et al., 2011; Kursawe et al., 2016; Sanchez-Gutierrez et al., 2013; Yamashita and Michiue, 2014). This approach has been adapted to analyze biomedical tissue samples, useful in clinical research and the development of diagnostic tools (Csikász-Nagy et al., 2013; Guillaud et al., 2010; Sáez et al., 2013; Sánchez-Gutiérrez et al., 2017). Finding features and patterns that can describe the graphs is key in many diverse fields, including biology (Benson et al., 2016; Costa et al., 2007; Hayes et al., 2013). A network can be split up into different subgraphs named graphlets. The graphlet composition of a network has been used to quantify differences between complex systems (Hayes et al., 2013; Ho et al., 2010; Kuchaiev et al., 2011; Pržulj et al., 2004). These measurements are based on the comparison of the quantity of each subgraph in different networks, providing an index of distance between them. This feature has the advantage of integrating the differences between diverse networks into a single value, simplifying the analyses and allowing multiple comparisons.

In summary, there is a clear need for a method to specifically quantify tissue organization and aid the interpretation of biophysical and mechanical aspects of morphogenesis and tissue homeostasis. The advances in imaging techniques, together with the appearance of powerful methods for automated image analysis (Heller et al., 2016; Khan et al., 2014; Kursawe et al., 2016; Schindelin et al., 2012; Weigert et al., 2018) and new simulation resources (Bi et al., 2016, 2015; Blanchard et al., 2009; Etournay et al., 2016; Fletcher et al., 2014; Guirao et al., 2015; Mirams et al., 2013; Tanaka et al., 2015) provide a large amount of good quality source data that can now be analysed in terms of organization. Here we present an open source platform, EpiGraph, a new image analysis method that uses segmented images from real epithelia or simulations, to easily quantify and compare the organization of packed tissues.

## RESULTS

### Graphlet measurements as an approach to capture organization of packed tissues

In previous studies, a set of 29 graphlets was used to distinguish between different types of networks (Pržulj et al., 2004) (**Fig. S1**). This method calculated the Graphlet degree Distribution agreement Distance (GDD) between two networks (Pržulj, 2007). Therefore, the “GDD value”, that in theory can range from 0 to 1, weighs the differences among the two distributions of graphlets; the higher the value, the more different the arrangements (**Fig. S1** and **Methods**). Epithelial images can be considered as natural tessellations and converted into networks of cell-to-cell contacts (Escudero et al., 2011). We have used the “graphlet” approach to capture the topology of epithelial tissues, making a correlation between graphlets and cellular motifs (compare Fig. 1A and **Fig. S1**). Tessellations give rise to “geographic networks” (Albert and Barabasi, 2002) that only make sense in a planar surface. For this reason, when we translated the set of graphlets to cellular patterns, some of them were redundant or not possible (see methods). Therefore, in this study we have used a total of 26 graphlets corresponding to 29 different cellular motifs that account for the organization of groups of up to 5 cells (Fig. 1A, **Fig. S1**). Most of the analyses performed in this work were completed with only 17 motifs (17-motifs, Fig. 1A, mauve). We found that, although all the motifs could be present in an actual tissue, 17-motifs minimized the redundancy of the information provided by the graphlets. In addition, this set downplays the importance of rare cellular geometries that could excessively weight GDD calculations (for example, a high difference in GDD could appear when comparing an image with one or two quadrilateral cells versus another image with no four-sided cells; this effect is minimized using 17-motifs). However, it would be possible to use other combinations such as all the motifs (29-motifs) or cellular motifs that account for the organization of groups of up to 4 cells (10-motifs) (Fig. 1A).

**Figure 1.**
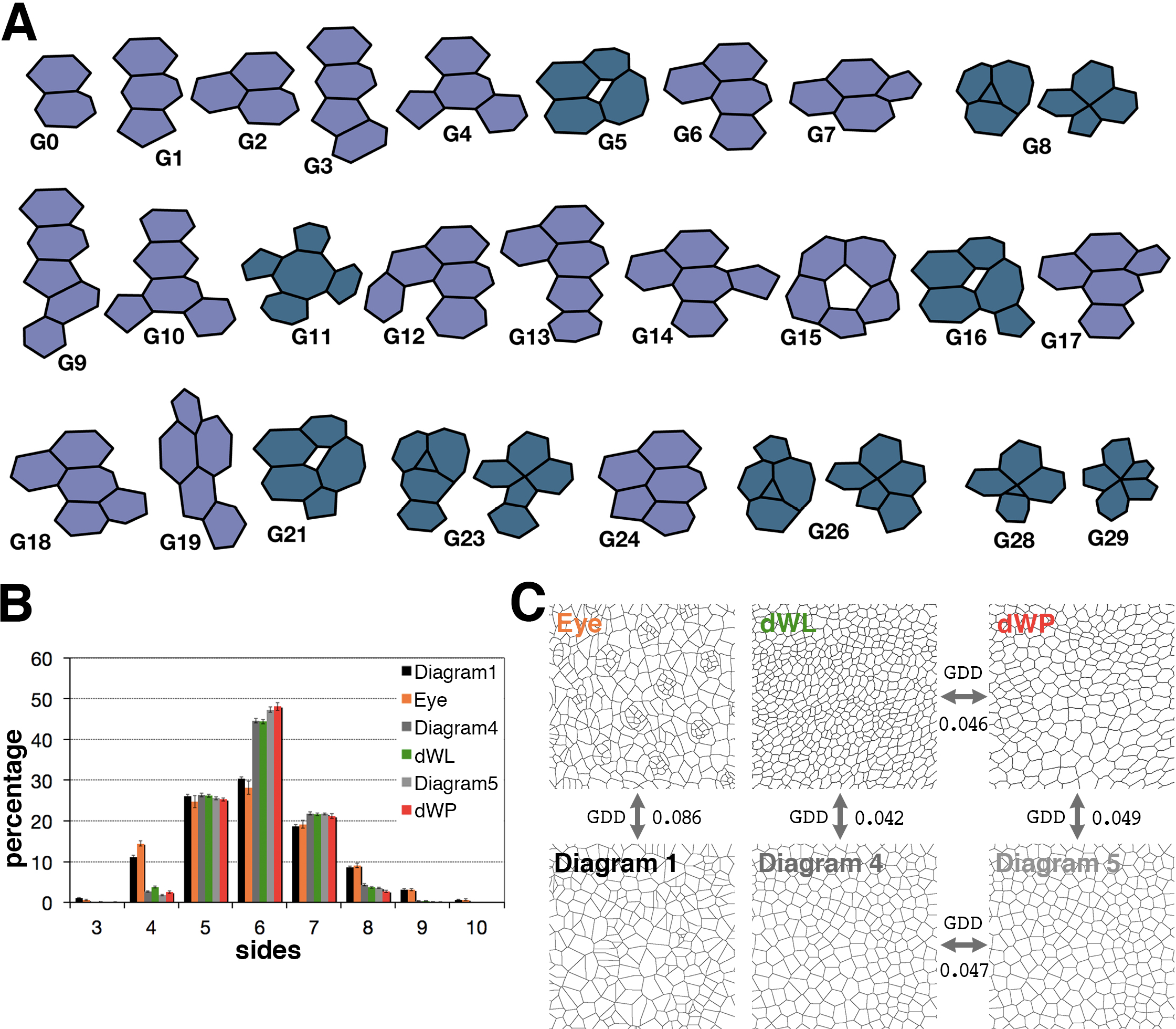
Graphlets, cellular motifs and characterization of epithelial organization. **A)** A representation of the cellular motifs that correspond to graphlets of up to five nodes. There are 29 motifs corresponding to 26 different graphlets (**Fig. S1**). Note that one graphlet can represent two cellular motifs (G8, G23 and G26). Mauve motifs form the 17-motifs set. Prussian Blue motifs stands for the set 29-motifs. In the first row are the motifs that account for the organization of groups of up to 4 cells (10-motifs). Therefore, 7-motifs set is formed by the mauve coloured graphlets at the first row. **B)** Polygon distribution comparison of images from: Voronoi diagram 1 (black bar); Eye (orange), *Drosophila* eye disc: 3 samples; Voronoi diagram 4 (grey); dWL (green), *Drosophila* larva wing disc: 15 samples; Voronoi diagram 5 (light grey), dWP (red), *Drosophila* prepupal wing imaginal disc epithelium: 16 samples. Data shown refer to the mean ± SEM. Diagram 1, 4 and 5: 20 replicates. **C)** GDD value calculation (17-motifs) between natural images and Voronoi diagrams with similar polygon distribution. The data shown are the mean of the GDD between each pair of images.

### Graphlet measurements capture differences beyond polygon distributions

We tested the power of graphlet-based measurements in quantifying differences between sets of images with very similar polygon distributions (Fig. 1B). In third instar larvae of *Drosophila*, the photoreceptors are specified, giving rise to a particular repetitive arrangement of the presumptive eye cells (Eye, Fig. 1C). This arrangement is very different to the irregular distribution in a Voronoi tessellation where the initial seeds were placed in a random way (Sanchez-Gutierrez et al., 2016), (Diagram 1, Fig. 1C). We previously showed that it was not possible to discriminate between the polygon distributions of these two tessellations (Sanchez-Gutierrez et al., 2016). Using the graphlets approach, we obtained a GDD value of 0.086 when comparing these two sets of images (17-motifs, **Table S1**, **Fig. S2**). In order to know if this difference was biologically relevant, we tried to set a baseline, by comparing other images with very similar polygon distribution that also presented an apparently similar arrangement. This was the case for Diagram 4 of the CVT vs. the *Drosophila* wing imaginal disc in larvae (dWL) and Diagram 5 of the CVT vs. the *Drosophila* wing imaginal disc in prepupae (dWP) (Fig. 1B-C). Both results were in the same range, with a GDD value of 0.042 for Diagram 4 vs. dWL and 0.049 for Diagram 5 vs. dWP (Fig. 1C). Similar results were obtained when comparing Diagram 4 vs. Diagram 5 and dWL vs. dWP (Fig. 1-C). These results suggested the existence of a baseline in the range of 0.04-0.05 values that correspond to similar cellular arrangements that cannot be well distinguished using the graphlets distribution. Therefore, we interpreted the value of 0.086 obtained in the Eye vs. Diagram 1 comparison as the reflection of actual differences between these two sets. In all the mentioned cases, the results obtained using 17-motifs and 29-motifs were equivalent (**Table S1**).

### EpiGraph quantitatively compares the organization of multiple sets of images

The GDD had the limitation of comparing only 2 samples each time. Here we have tried to overcome this limitation evaluating different images simultaneously using a reference. Therefore, we designed EpiGraph, a method that calculates the GDD of any epithelial tissue with another tessellation that serves as a reference. We used three different references: i) a tessellation formed by regular hexagons, representing the most ordered way to pave the space (Fig. 2A, Epi-Hexagons). ii) the network motifs emerging from a random Voronoi tessellation (Fig. 2B, Epi-Random). iii) a Voronoi Diagram 5 from the CVT path (Fig. 2C, Epi-Voronoi5) that presents a polygon distribution similar to the one from multiple examples in nature (Gibson et al., 2006; Sanchez-Gutierrez et al., 2016).

**Figure 2.**
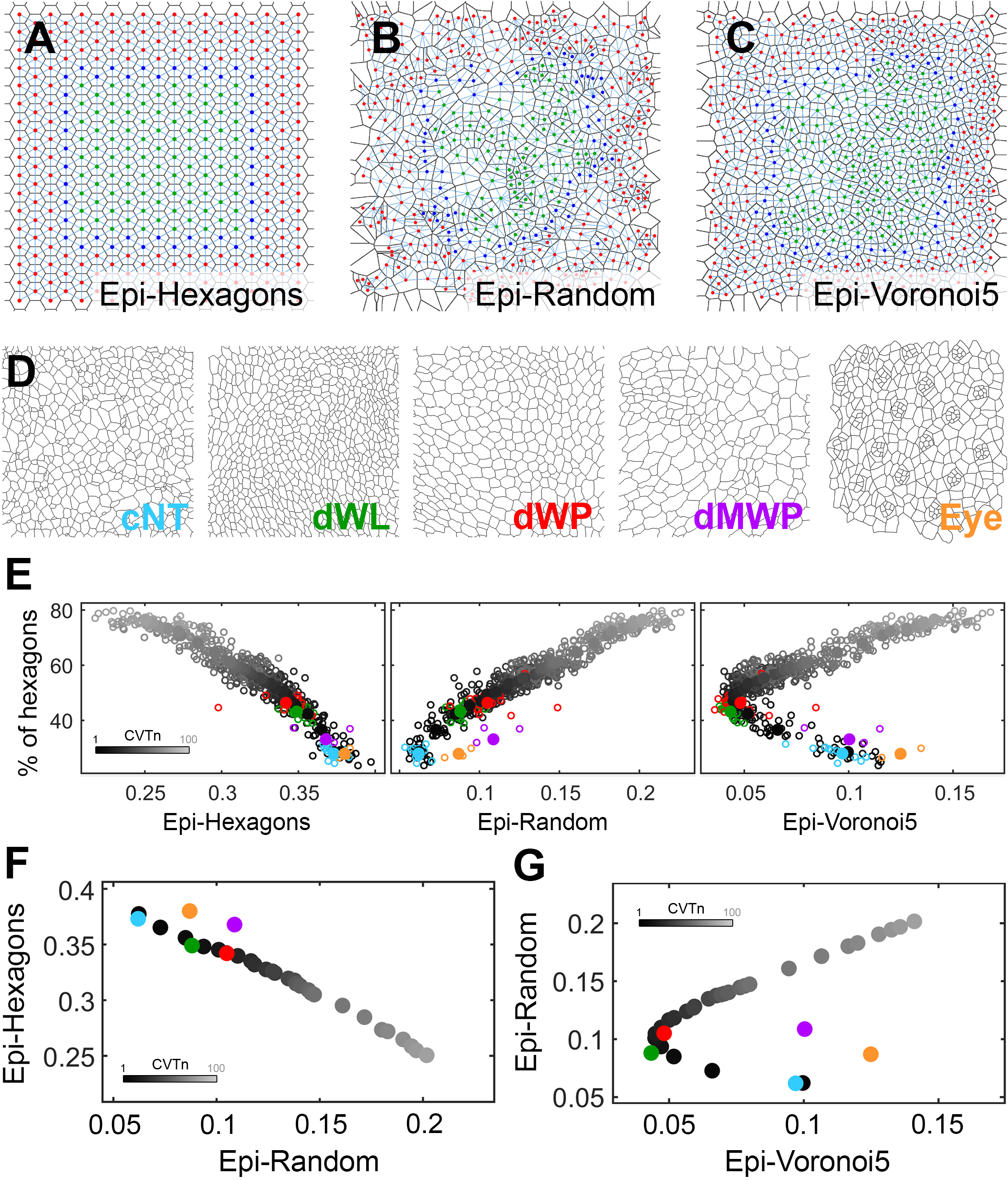
Epithelial organization of biological tissues with respect to the CVTn. **A-C)** Tessellations with the corresponding graph of cell-to-cell contacts for a perfect hexagonal arrangement (**A**) a Voronoi Diagram 1 (**B**) and a Voronoi Diagram 5 (**C**) from a CVTn. These tessellations represent the diagrams used as reference to calculate the Epi-Hexagons, Epi-Random and Epi-Voronoi5 respectively. The light blue edges in these panels represent the cellular connectivity network. The colourful nodes mark the valid cells that were involved in the cellular motifs to measure graphlets presence. The dark blue and green nodes are the 3-distance valid cells (cells connected exclusively to valid cells within a distance of 3 edges), which were used to calculate the graphlets for 10-motifs and 7-motifs. The green nodes are the 4-distance valid cells (cells connected exclusively to valid cells within a distance of 4 edges) that were used to quantify the graphlets for 29-motifs and 17-motifs. Cells without nodes were no valid cells for graphlet calculation. **D)** Representative images from the natural tessellations. **E)** Plots showing the different combinations of the values for 17-motifs of Epi-Hexagons, Epi-Random and Epi-Voronoi5 with the percentage of hexagons. The diagrams of the CVTn path from the iteration 1 until the iteration 700 are represented as a greyscale beginning in black and reducing its darkness with the increase of the iterations (from 1 to 20, from 30 to 100 by stepwise of 10 and from 100 to 700 by stepwise of 100). **F-G**) Charts representing the comparisons Epi-Hexagons against Epi-Random, and Epi-Random against Epi-Voronoi5, respectively. The CVTn path in both scatter plots, is formed by the diagrams with numbers between 1 and 100, in a greyscale as in (**E**). The natural tessellations are: dMWP (violet), *Drosophila* mutant wing disc: 3 samples; cNT (light blue), chicken neural tube: 16 samples; Eye, dWL and dWP are the same replicates than Fig.1 and preserve their colour codes. Circumferences are individual values, circles are the average value obtained from the individual samples from each category.

We tested the method with epithelial images that have been previously compared with the CVT path in terms of polygon distribution: chicken neural tube (cNT), dWL, dWP, reduction of myosin II in the *Drosophila* prepupa wing disc epithelium (dMWP) and Eye (Fig. 2D)(Sanchez-Gutierrez et al., 2016). To have a scale and facilitate fast comparisons, we used the concept of the CVT path (Sanchez-Gutierrez et al., 2016). We calculated the GDD values for Epi-Hexagons, Epi-Random and Epi-Voronoi5 for all the Voronoi diagrams and visualized these results with respect to the percentage of hexagons of the corresponding diagram (the percentage of hexagons is indicative of the proportions of the different types of polygons along the CVT, **Table S2**). However, the CVT does not progress beyond the 70% of hexagons limiting the possibilities of analysis. Therefore, we extended the Voronoi scale spanning a wider range of polygon distributions. The algorithm that devises the CVT was modified to introduce “noise” in the positioning of the seed that produces the subsequent diagram. In this way, we obtained a “CVT noise” (CVTn) whose last diagrams reached 90% of hexagons (Fig. 2E-G, **Fig. S3** and **Material and methods**). Interestingly, the plot obtained using CVT and CVTn diagrams was an optimum way to easily visualize these geometric scales as a continuous “CVT path” and a “CVTn path”. Therefore, we used this framework to analyse the values of Epi-Hexagons, Epi-Random and Epi-Voronoi5 for each diagram in the scale. As expected, the Epi-Hexagons values were higher in the initial diagrams and progressively decreased with the increase in the percentage of hexagons of the Voronoi diagrams (Fig. 2E, **left panel**). The opposite happened in the case of the Epi-Random values (Fig. 2E, **central panel**). In the plot of percentage of hexagons vs Epi-Voronoi5, the CVTn path presented the shape of a walking stick (Fig. 2E, **right panel**). The Epi-Voronoi5 values of Voronoi Diagrams 1, 2, 3, and 4 were decreasing progressively, with Diagram 5 the closest to the zero value. The values for the rest of the diagrams gradually increased, as in the case of the Epi-Random.

We then plotted the values for the actual epithelia. We found that for cNT, dWL and dWP the Epi-Hexagons, Epi-Random and Epi-Voronoi5 values were similar to the CVTn at the same percentage of hexagons of the polygon distribution (Fig. 2E). In agreement with our previous results using the GDD, the Eye images presented a higher Epi-Random and Epi-Voronoi5 values than the expected for a 30% of hexagons (Fig. 2E). The differences with respect to the CVTn were even more clear when plotting Epi-Hexagons vs Epi-Random and Epi-Random vs Epi-Voronoi5 (Fig. 2F-G). We obtained similar results when analysed the dMWP set of images. In this case, our previous work showed a small deviation of the dMWP polygon distribution with respect the CVT (Sanchez-Gutierrez et al., 2016). However, using Epigraph, we observed that Epi-Random and Epi-Voronoi5 captured the clear differences in organization between these images and the CVTn (Fig. 2E-G, **Fig. S4**). These results suggested that EpiGraph is able to distinguish between different tessellations with a similar polygon distribution. In this regard, we have developed a statistical output using an outlier detection approach whose quantitative results represent how similar the organization of a tissue is when compared with the CVTn scale (**Fig. S3** and **Material and methods**). The test confirmed that cNT, dWL, and dWP were close to the CVTn and similar to the Voronoi diagrams 1, 4, and 6 respectively. In contrast, the Eye and dMWP samples were labelled as different (**Table S3**). In this way, EpiGraph provides a quantitative description of tissue organization.

### Epigraph can capture different organization traits

We further investigated the possible applications of EpiGraph and performed a series of experiments aimed at understanding what traits of tissue organization are being captured and quantified by the graphlet measurements. To this end, we have used images of different vertex model simulations that alter tissue organization by changing the biophysical properties of the cells (images taken from (Sanchez-Gutierrez et al., 2016) (**Material and methods** and Fig. 3A-F).

**Figure 3.**
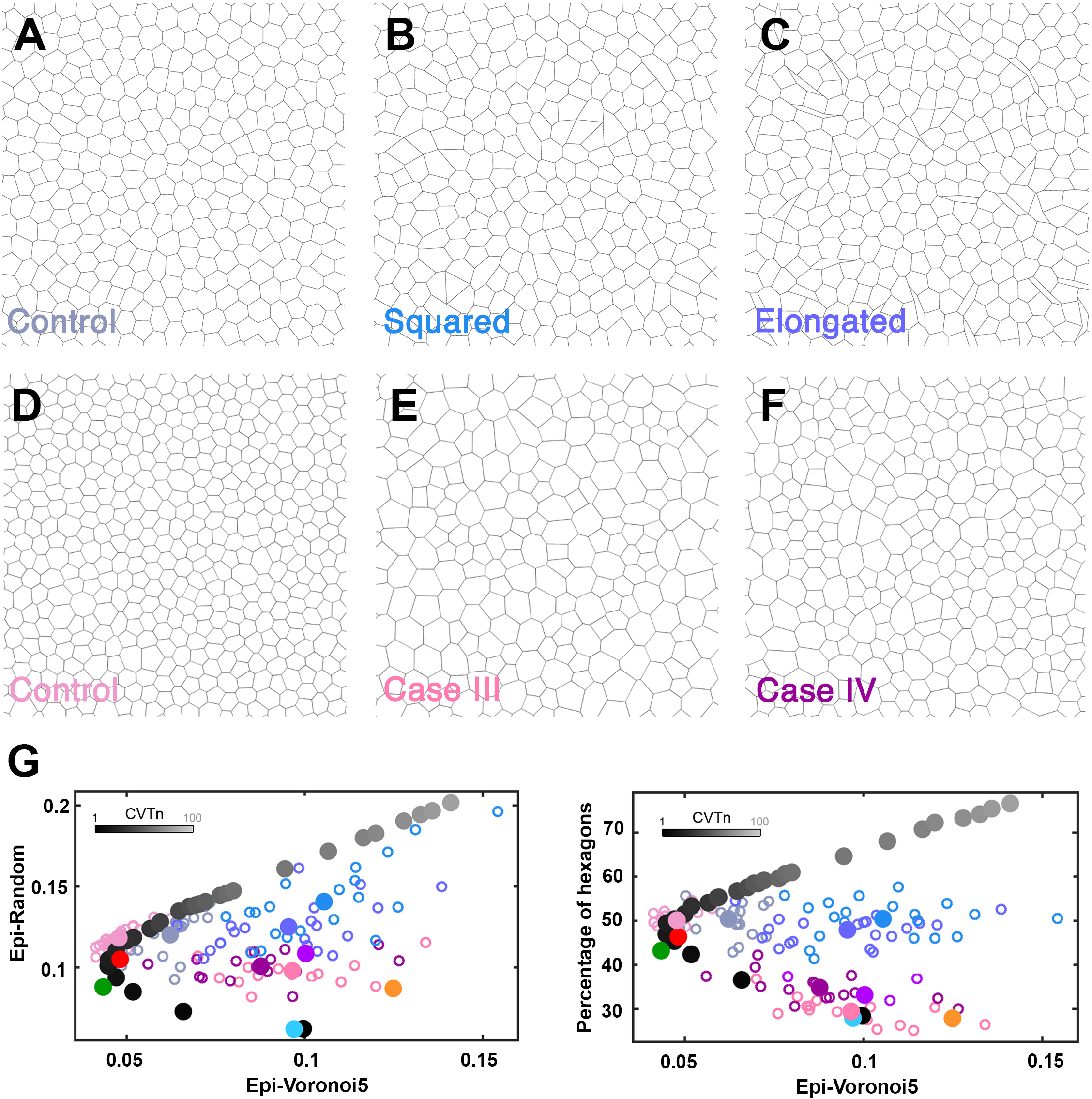
Comparison of different simulations and mutants with the CVTn. **A-C)** Representative images for non-proliferative simulations. Control with homogeneous parameters (**A**). The ‘squared’ simulations are similar to control, but a ten percent of cells (randomly chosen) have a reduced line tension (**B**). The ‘elongated’ simulations have a ten percent of cells (randomly chosen) with its line tension and ideal area reduced, and the another 90% of cells have the same parameters than control simulations (**C**). **D)** Cell arrangement resulting from the control simulation that includes cell proliferation. **E-F)** Diagrams resulting from a vertex model simulation with an increase of the ideal area value, with respect the control, in some cells. Case III and Case IV slightly differ in the line-tension parameter conditions (see **Material and methods**). **G)** Plots showing the values of Epi-Random vs Epi-Voronoi5 and the percentage of hexagons vs Epi-Voronoi5 (17-motifs) for CVTn, dMWP, Eye, cNT, dWL, dWP; Proliferative Control (20 replicates, carnation pink), Case III (17 replicates, hot pink) and Case IV (15 replicates, purple); Non-proliferative control (20 replicates, blue bell), Squared (20 replicates, azure blue) and Elongated simulations (20 replicates, cornflower Blue). The diagrams of the CVTn path from the iteration 1 until the iteration 100 are represented as a greyscale beginning in black and reducing its darkness with the increase of the iterations; dMWP, Eye, cNT, dWL and dWP have the same replicates and colour codes than Fig.2; circumferences are individual values, circles are the average value obtained from the individual samples from each category.

First, we analysed samples with 10% of the cells having increased effective cell-cell adhesion (**Material and methods**, Fig. 3B). This feature induced the formation of cells with a “quadrilateral shape” that often organized in motifs presenting four-way vertex configurations. These images were compared with simulations in which “elongated” cells appear (by simultaneously increasing cell-cell adhesion and reducing ideal area, **Material and methods** and Fig. 3C). Epigraph analysis indicated that while control simulations gave similar values to the CVTn, the “squared” and “elongated” sets of images were different to the control and well separated from the CVTn. However, EpiGraph failed to find clear differences between the “squared” and “elongated” images (Fig. 3G and **Fig. S5**).

Second, we used a set of conditions to mimic the effect of a reduction of myosin II in the *Drosophila* prepupa wing disc epithelium (dMWP, Fig. 2D). In the control simulation (Fig. 3D), cells grow to double the original area and then divide into two cells. In case III and case IV simulations there was a random reduction of the tension parameter together with a requirement of a minimum tension threshold to be able to divide (Fig. 3E-F). If the cells do not reach this threshold, they continue to grow without dividing the cell body. When this happens, the cells will be stuck in mitotic phase and will not start a second round of cell division (Sanchez-Gutierrez et al., 2016) (**Material and methods**). The control simulation gave similar values to the CVTn, while case III, case IV and dMWP images presented a clear deviation in the Epi-Random vs Epi-Voronoi5 graph (Fig. 3G). All these data-points distributed in the same zone of the graph. Interestingly, we found that both sets of simulations (squared and elongated vs Case III and Case IV) appeared in two complementary regions, suggesting that the regions in the graph can reflect the existence of different traits of organization in each condition (Fig. 3G).

### EpiGraph: a method to capture epithelial organization implemented in FIJI

Aiming to enhance the accessibility of the analysis of tissue organization to the biology community, we have implemented EpiGraph as a plugin for FIJI (Schindelin et al., 2012). EpiGraph consists of a pipeline of 5 very simple steps. First, the skeleton of an epithelial image is uploaded and the individual cells are identified. Second, the user selects the distance threshold to identify two cells as neighbours. Here it is possible to select different thresholds and to check the number of neighbours of every cell in each case. Third, a ROI is selected. There are several possibilities such as a default ROI from the image or the selection of individual cells. Fourth, the graphlet information for the selected cells is calculated. These data are used to obtain the Epi-Hexagons, Epi-Random and Epi-Voronoi5. These values are incorporated into a table and serve as input data for a statistical analysis that indicates if a new image is inside or outside of the CVTn path and describes which Voronoi diagram presents the most similar organization to the sample (**Material and methods**). The fifth step includes the classification and labelling of different images in order to represent them in a new window. This final phase allows one to export the representation of the data in a three-dimensional graph. **Movie S1** shows an example of EpiGraph usage. A detailed description of EpiGraph can be found in the **Supplementary Material and methods**. A full set of tutorials explaining how to install and use EpiGraph is available at EpiGraph’s wiki (https://imagej.net/EpiGraph).

### EpiGraph provides biological insights regarding homeostasis and tissue fluidity transitions

Epithelial tissues have the ability to behave as a fluid due to cellular rearrangements or to solidify as cellular rearrangements cease (Bi et al., 2016, 2015). The shape index is a characteristic of epithelial cells that has been shown, in vertex model simulations, to be able to capture the degree of rigidity, or fluidity, of a tissue (Bi et al., 2015). This study established the transition point between a soft (fluid) and a rigid (solid) tissue, described as a jamming transition, at the dimensionless shape index value of 3.81. We calculated the shape index for the CVTn path, finding that from Voronoi diagrams 1 to 20, the tessellations were behaving as a fluid (from diagram 21 to 700 they behave as solid). Using this descriptor, all the images of biological tissues were placed in the fluid part as well as the four altered vertex model simulations shown in Fig. 3 (**Fig. S4**, **Fig. S5**, **Fig. S6** and **Table S4**).

We have investigated the dynamics of epithelial jamming in different conditions. First, to test the capabilities of EpiGraph in this regard, we analysed several snapshots from two simulations published by Bi and colleagues as supplementary movies (Bi et al., 2016). These videos show the movements of cells in two conditions: rigid state (shape index less than 3.81) and soft state (shape index greater than 3.81) (Fig. 4A). As expected, the snapshots of the soft tissue analysed appeared in different positions, indicating that the simulated epithelia changed its organization during the experiment. On the other hand, the different frames from the rigid simulation were clustered (Fig. 4B), showing little cell rearrangements.

**Figure 4.**
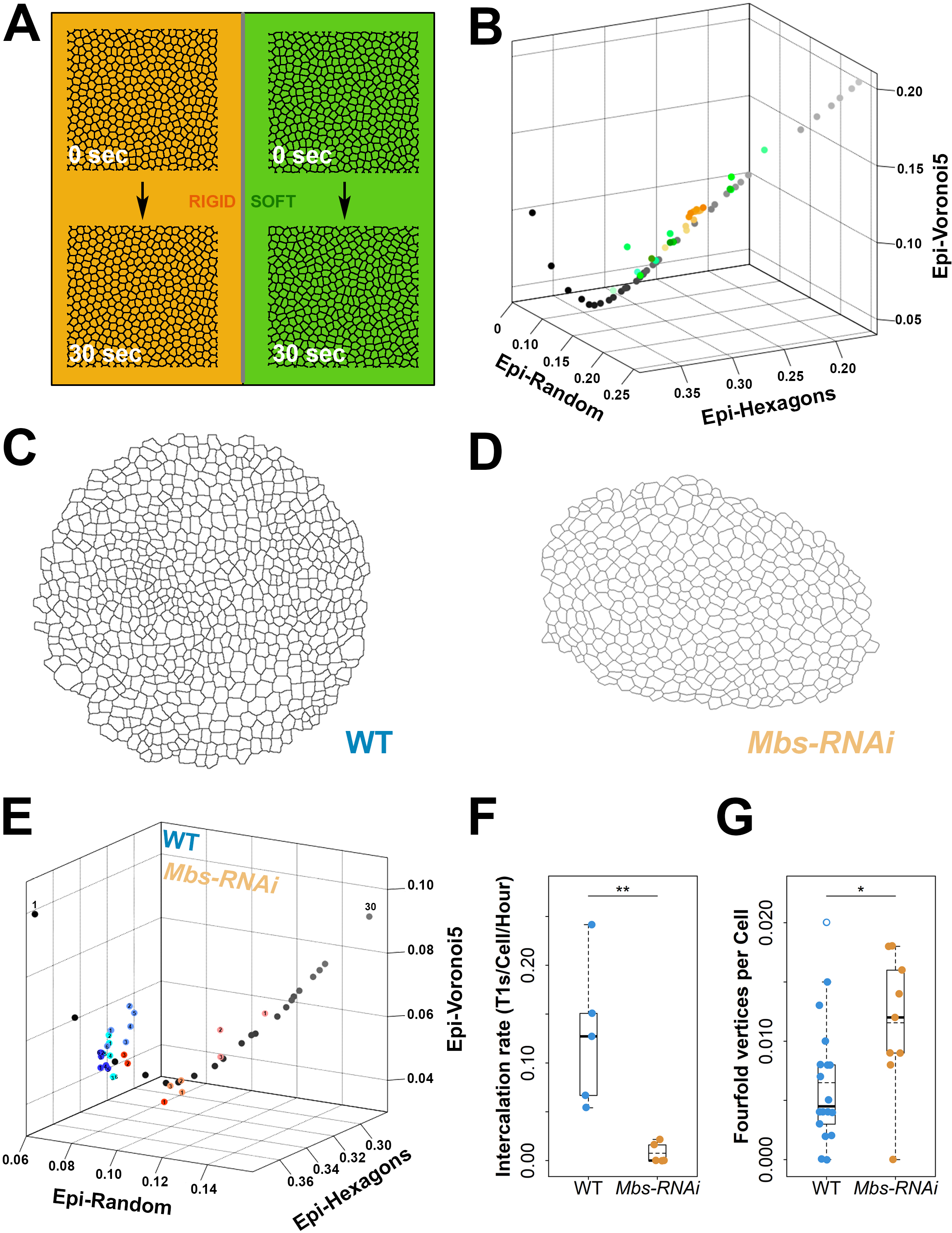
The rigidity/fluidity of a tissue can be assessed using EpiGraph. **A**) Initial and final frames of two simulations with different settings: a rigid and a soft tissue. **B)** Epigraph’s 3D plot with Epi-Random, Epi-Hexagons and Epi-Voronoi5 axes, showing the soft simulation tissue in green dots and the rigid simulation as orange dots. Each simulation is represented in 13 frames (see **Material and methods**). **C-D)** Representative examples of segmented images from the third instar *Drosophila* imaginal disc in different conditions: Wild Type (**C**); a solid mutant, *Mbs-RNAi* in (**D**). **E)** Plot comparing the fluidity and organization of the tissues in (**C-D**). CVTn (until diagram 30) displayed in greyscale. Dots in scales of blue represent the WT condition: wing disc 1, aquamarine; wing disc 2, light blue; wing disc 3, dark blue. Represented with points in tones of orange-red, *Mbs-RNAi*: sample 1, salmon colour; sample 2, orange; sample 3, red. **F-G)** Boxplot of the intercalation rate (**F**), which is the number of T1 transitions per cell per hour, and the fourfold vertices found per cell (**G**) (note that no fivefold vertices, or beyond, was found on any sample). Boxes stand for the data inside the upper and lower quartiles, while the vertical dashed lines (whiskers) indicate the variability outside them. Mean (dashed line) and median (thick line) of each condition is represented inside each box. The actual values are also presented as circles (and the outlier values as circumferences) with its correspondent colour. In addition, statistical significance, by means of a Kolmogorov-Smirnov test, is shown in the top of both panels (**F**: ‘**’ p<0.01, **G**: ‘*’ p<0.05). Each condition has 3 samples (different colour tone), and the numeric tag represents its frame. In WT have been taken 6 frames per sample in periods of 6 minutes. In the case of *Mbs-RNAi* were taken 3 frames per sample with time lapse of 15 minutes. All the conditions have been tracked for 30 minutes (see **Material and methods**).

We next tested whether EpiGraph could detect changes in tissue fluidity in real epithelia, which may be more ambiguous and noisier than simulations. Real tissues also display fluid-to-solid jamming transitions which are important for large scale tissue shape changes as well as for refining and maintaining tissue shape (Curran et al., 2017; Mongera et al., 2018). In the *Drosophila* pupal notum, the level of tissue fluidity is controlled by the global level of myosin II activity (Curran et al., 2017). We wondered if the regulation of myosin II could similarly impact on the fluidity state of the wing disc epithelium and the cell rearrangements that have been described during the late stages of normal wing disc development, where the overall tissue shape does not dramatically change (Heller et al., 2016) (Fig. 4C). To this end, we compared the WT organization with the effect of increasing myosin II activity by knocking down Mbs (Myosin binding subunit of the myosin phosphatase, which dephosphorylates myosin regulatory light chain, Fig. 4D) by RNAi throughout the entire wing pouch. Based on work in the pupal notum (Curran et al., 2017), we would predict that *Mbs-RNAi* discs behave as solids. Interestingly, in the two cases, the shape index was greater than the described shape index threshold of 3.81, suggesting that both tissues are in a fluid state (**Fig. S6**).

We used EpiGraph to analyse the changes in organization of wing discs with perturbed myosin II activity along time and compared them with a WT condition. The snapshots for WT samples appeared clustered in the 3D graph, indicating that the epithelia were not changing their organization during the 30 minutes of analysis (Fig. 4E), despite previous work showing that cell intercalations do occur (Heller et al., 2016). This suggests that during this slow growing phase of wing disc development, any cell rearrangements that occur do not drive large-scale morphogenesis, but act to maintain a homeostatic tissue topology. The statistical analysis confirmed that all of the WT wing discs were close to CVTn diagrams 3 and 4 (**Table S3**). In the case of the three samples from the *Mbs-RNAi* genotype, the data points presented different organizations (from similar to diagram 3 to close to diagram 13, see **Table S3**). In some cases, the dispersion was not only between samples, but occurred between images from each movie (Fig. 4E and **Table S3**). EpiGraph therefore predicted that these *Mbs-RNAi* wing discs are behaving very differently from WT wing discs, likely by changing their degree of fluidity. Accordingly, quantification of intercalation rates demonstrated that cell rearrangements happen significantly more frequently in WT than in *Mbs-RNAi* wing discs (Fig. 4F, 0.1281±0.08 vs. 0.0076±0.01 intercalations/cell/hour, Kolmogorov-Smirnov test, *p*=0.0079). As predicted, this resulted in more cells ‘jamming’ at 4-way vertex configurations as they fail to complete intercalations (Fig. 4G, 0.0065±0.005 vs. 0.0116±0.006 fourfold vertices/cell, Kolmogorov-Smirnov test, *p*=0.029). Interestingly, EpiGraph was able to detect this solidification of the tissue in the *Mbs-RNAi* discs, even though the shape index predicted a fluid tissue. Taken together, these results indicate that the quantification of tissue organization using EpiGraph can infer information about the fluidity of a tissue from several fixed snapshots, without the need to laboriously track individual frames of a time-lapse video.

## DISCUSSION

Textbook definitions of morphogenesis include the term “organization” as key to explaining this fundamental developmental process (Dai and Gilbert, 1991). The authors wondered, “How can matter organize itself so as to create a complex structure such as a limb or an eye?”. Later, changes in organization of adult tissues can reflect pathological traits due to defects in homeostasis (Csikász-Nagy et al., 2013; Soto and Sonnenschein, 2011). Here, we have provided a tool that can help to investigate these questions.

The analysis of the polygon sides of epithelial cells has been shown to be insufficient to completely understand tissue organization. Some tessellations can present very different arrangements yet have the same frequencies of number of neighbours. A second problem is the lack of a simple value as an indicator of epithelial organization. This feature complicates the comparison between morphogenesis of normal development and that of genetically perturbed or diseased tissues. Our previous attempts to overcome this caveat were based on multi-statistical analyses of graph features (Sanchez-Gutierrez et al., 2013) and the creation of a Voronoi scale to statistically compare groups of images with the CVT reference (Sanchez-Gutierrez et al., 2016). Several recent works cover part of these integrative analyses (Blanchard, 2017; Blanchard et al., 2009; Farrell et al., 2017; Guirao et al., 2015; Jackson et al., 2017). However, we are aware that all these methods are difficult to incorporate into the average biology or biomedicine lab.

We have developed EpiGraph, aiming to bring an easy way to quantify tissue organization without the requirement for programming skills. EpiGraph transforms the image into a graph of cell-to-cell contacts and extracts their graphlet content to later compare with other images. These complex algorithms are hidden behind the friendly user window of FIJI. This is the most popular open-source biological image analysis platform. In addition, the output data options of EpiGraph facilitate fast and clear representations and interpretations of the results.

One of the strengths of EpiGraph is the comparison of any tessellation with the hexagonal lattice, the “random” Voronoi tessellation and the Voronoi tessellation that presents the “conserved polygon distribution” (Gibson et al., 2006; Sanchez-Gutierrez et al., 2016) (Fig. 2A-C). We have tested EpiGraph with different types of samples: as expected, the average of the natural tessellations such as wing imaginal disc (dWL and dWP) or the chicken neural tube (cNT) matched the CVTn path position (Fig. 2D-G). We interpret that these three natural samples present similar polygon distributions and graphlet compositions to some Voronoi Diagrams from the CVTn. On the other hand, the average of the Eye samples appeared far from the CVTn when Epi-Voronoi5 or Epi-Random values were plotted (Fig. 2D-G). These two references were capturing differences in organization between the Eye and any Voronoi Diagram (including Diagram 1, which presents a similar polygon distribution to the Eye). This result supports the utility of EpiGraph to quantify organizational traits that were not accessible until now. The same idea is reinforced by the results obtained with the mutant samples for myosin II (dMWP, Fig. 2D-G). In previous work, we showed that this set of samples slightly deviated the CVT scale in terms of polygon distribution (Sanchez-Gutierrez et al., 2016). Here we show very clear differences in terms of the values of Epi-Voronoi5 and Epi-Random (Fig. 2G), suggesting a higher sensitivity of the new method when capturing differences in organization.

The output images from EpiGraph show the CVTn path as a clear reference for proliferative epithelia such the wing imaginal disc or the chicken neural tube and for vertex model control simulations. We have incorporated a statistical test into EpiGraph that indicates if a new tissue is within or outside of the CVTn path, and which is the Voronoi diagram with the closest organization. The different results comparing Epi-Hexagons, Epi-Random and Epi-Voronoi5 values also suggested that Epi-Hexagons had better resolution for images with a higher percentage of hexagons while Epi-Random and Epi-Voronoi5 were more sensitive to the differences between images with less than 40% of hexagons. For this reason, we have designed the visualization step of the program to easily change the three axes and check the different results using any combination of these GDD references and the “percentage of hexagons”.

Using different sets of simulations, we are able to distinguish two different types of organization: The cases where a subset of cells adopts a particular arrangement inside a mostly ordered tissue (Fig 3B, C, G) and the cases where the global topology of the tissue is altered and the cell sizes are very heterogeneous (Fig. 3E-G and **Fig. S5**). These two patterns create a “map” of arrangements that are out the CVTn, and they will help to other researchers to study the degree of order in their samples.

The dynamics of the transition between a tissue behaving as a fluid or a solid is an emerging problem in developmental biology and biomedicine (Curran et al., 2017; Firmino et al., 2016; Mongera et al., 2018; Park et al., 2015; Petridou et al., 2018; Tetley and Mao, 2018). We have used the capabilities of EpiGraph to study how the fluidity state can affect the organization of a tissue. The utility of EpiGraph in this regard is supported by its ability to quantify dynamic changes in organization due to cell rearrangements in a vertex model simulation of a soft tissue (Fig. 4A-B). Therefore, in these simulations, cell movements are captured as changes in the organization of the tissue by EpiGraph. However, cell rearrangements do not necessarily have to lead to changes in tissue organization, as is often the case in more homeostatic tissues. Although, it has been shown that the late third instar *Drosophila* imaginal disc can exchange neighbours and rearrange during development (Heller et al., 2016), we were not able to see changes in organization combining live imaging of the WT discs and EpiGraph analysis (Fig. 4E). Therefore, we interpret that the multiple re-arrangements of the WT disc conserve the organization of the tissue, at least in the time framework analysed (30 min). On the contrary, the hyperactivation of myosin II (*Mbs-RNAi*) produced a clear change in the organization of the tissue as detected by EpiGraph. The decrease of intercalation rate and the increase of fourfold vertices in the *Mbs-RNAi* discs suggest that EpiGraph is capturing a change in tissue fluidity (Fig. 4E-G). In this respect, we think that EpiGraph analyses provide information beyond previous parameters that have been used to capture the fluidity in cell arrangements such as the shape index (Bi et al., 2015). All the real images analysed in this work have a high shape index (**Fig. S6**). These samples include the *Mbs-RNAi* mutant discs, that do not intercalate. Altogether, our results suggest that the shape index is not a sufficient parameter to define fluidity from a still image of a real sample.

In biomedicine, a robust and efficient analysis of histopathological images is required. Computerized image tools have an enormous potential to improve the quality of histological image interpretation, offering objective analyses that can aid the pathologist’s diagnoses. Changes in organization have proven to be related to the onset of disease in very different contexts, being critical for early detection (Emmanuele et al., 2015; Guillaud et al., 2010; Park et al., 2015; Sáez et al., 2013; Tsuboi et al., 2018). We propose that EpiGraph is able to efficiently detect mutant phenotypes related to changes in organization and/or in tissue fluidity. Importantly, this can be done from a few snapshots in time, without the need for sophisticated time-lapse imaging and tracking. This may provide a simple detection tool for the early onset of disease, where changes in organization can occur, and only limited tissue samples are available from patients.

### EpiGraph limitations

Although Epigraph accepts a wide range of images as inputs, we have specified some minimum requirements. It is not recommended to use input images bigger than 3000 pixels of width or 3000 pixels of height, since processing them could be computationally intensive. In addition, EpiGraph only accepts single images. Images from time series should be adapted to single frames before uploading them to EpiGraph.

Computers with little RAM memory (less than 16gb) will work but with a series of restrictions. To ensure usability, it is not recommended computing images with a high number of cells (more than 1000) due to a possible lack of memory. In the same way, we suggest skeletonizing the edges of the images and using a small radius, i.e. 3 pixels of radius for skeletonized image (we recommend don’t overpass a radius value of 10 pixels to avoid overloading the system) to calculate the cells neighbourhood. Choosing a high radius value could slow down the work queue, increasing the use of RAM memory.

If any of these requirements are not satisfied, the program alerts the user, allowing him/her to change the image provided. Importantly, the images and ROIs require a minimum number of valid cells (cells without touching the borders or an invalid region of the image) in order to get coherent graphlets. Therefore, to get any result, EpiGraph must detect at least a 3-distance valid cell (see Fig. 2) in the case of 7-motifs or 10-motifs or a 4-distance valid cell (see Fig. 2) in the case of 17-motifs and 29-motifs. In any case, we strongly recommend having a greater number of 3-distance and 4-distance valid cells to get results that can be trusted in terms of capturing the organization of a tissue. Regarding the 3D visualization tool, it allows the user to see the position of the samples from different angles. However, the resolution of the exported file is only 72 pixels per inch (dpi). This could be too low for publications and therefore EpiGraph provides an excel table with all the information needed to represent it with other programs.

In summary, we have generated a very accessible, open source method to produce a quantitative description of developmental events. This quantitative aspect is reinforced by the statistical comparison with the CVT path that serves as a scale for tissue organization. We anticipate that our tool will improve the study of tissue dynamics and morphogenesis by permitting the comparative analysis of epithelial organization in genetically mutated or diseased tissues during time.

## MATERIAL AND METHODS

### Source images used in the study

#### Centroidal Voronoi Tessellation (CVT) diagrams and variations

For the generation of this set of paths we have used the software Matlab R2014b to iteratively apply Lloyd’s algorithm to a random Voronoi tessellation (Lloyd, 1957). This implies that the centroid of a cell in a Voronoi diagram is the seed for the same cell in the next iteration.

##### Centroidal Voronoi Tessellation (CVT) diagrams

Centroidal Voronoi Tessellation diagrams were obtained as described previously by our group (Sanchez-Gutierrez et al., 2016). The 20 original Voronoi diagrams were created placing 500 seeds randomly in an image of 1024×1024 pixels. A total of 700 iterations were generated for each initial image.

##### Centroidal Voronoi Tessellation noise (CVTn) diagrams

We have developed a variation of the CVT path, named the CVT noise (CVTn) path (**Fig. S3**). We started from the same 20 initial random diagrams described above. The development process of the CVTn path was modified so that the new seeds were not strictly the centroid from the previous iteration. In even iterations, we selected a region of 5 pixels of radius from the centroid position, in which seeds could be placed randomly. In odd iterations, the system was stabilized, applying the original Lloyd algorithm. A total of 700 iterations were generated for each initial image.

#### Natural packed tissues and vertex model simulations

The details of the obtaining and processing of the epithelial images were described in (Escudero et al., 2011). Control vertex model simulations include cell proliferation and are the basis for the other two cases. Case III corresponds to a vertex model simulation with heterogeneous reduction of line tension and an impairment of cell division when tension value is under 30 percent of the initial value. Case IV is a similar simulation to Case III with a threshold of 40 percent. Regarding simulations with no cell proliferation, as a baseline, the control had homogeneous parameters for contractility, line tension and ideal area. ‘Elongated’ simulations were as the control, but with ten percent of cells having a reduced line tension and ideal area, while ‘squared’ ones had ten percent of cells with only line tension reduced. The exact conditions for the vertex model simulations were described in (Sanchez-Gutierrez et al., 2016).

#### Perturbing myosin II activity in *Drosophila* wing discs and calculating intercalation rates

*Drosophila* were raised in standard conditions. Wing discs were dissected from third instar larvae and cultured under filters as described by (Zartman et al., 2013). Discs were cultured in Shields and Sang M3 media supplemented with 2% FBS, 1% pen/strep, 3ng/ml ecdysone and 2ng/ml insulin. The following alleles and transgenes were used; *shg*-GFP (Ecad-GFP, Huang et al., 2009), *UAS-Mbs-RNAi* (KK library, VDRC), *rn*-GAL4 (RMCE-MiMIC Trojan-GAL4 collection). The following experimental genotypes were used; Ecad-GFP (WT) and Ecad-GFP/UAS-*Mbs*-*RNAi*; *rn*-GAL4/+ (*Mbs-RNAi*). For EpiGraph analysis, discs were imaged on a Zeiss LSM 880 microscope with Airyscan at 512×512 resolution with a 63x objective (NA 1.4) at 1.4x zoom for a total of 30 minutes with 1-minute time intervals and a z-step of 0.5µm. Time-lapse image sequences were segmented using Epitools (Heller et al., 2016).

To quantify intercalation rates, 5 WT and 5 *Mbs-RNAi* wing discs were imaged using the same methods as above, except using 5× zoom and 3 minutes intervals for a total of 2 hours. Intercalation rate data was exported from the “EDGE_T1_TRANSITIONS” overlay in the “CellOverlay” plugin in Epitools. To exclude mistakes generated when 4-way junctions were not recognised, junctions less than 0.075µm in length were assigned a length of 0µm. A productive intercalation event was scored when a neighbour exchange was stabilised for at least 2 time points (6 minutes). The total number of tracked cells was also quantified, allowing the intercalation rate to be expressed as the number of intercalations per cell per hour.

We also counted the number of fourfold vertices per cell in both WT and *Mbs-RNAi* conditions. In particular, we quantified the number of vertices in which four or more cells were touching each other, using Matlab R2014b. The cells closest to the border of the image were excluded from the analysis. In this way, we obtained the percentage of fourfold vertices per valid cell for each image and calculated a Kolmogorov-Smirnov test to check if the distributions of both conditions were different.

### Soft and Rigid tissue simulations

We have extracted a set of screenshots from two videos that simulated different dynamical behaviour of vertex model simulations. These videos are presented as Supplemental Material in (Bi et al., 2016). The first video represents a rigid behaviour in the simulation: https://journals.aps.org/prx/supplemental/10.1103/PhysRevX.6.021011/solid_tissue_v0_0.2_p0_3.5_Dr_0.1.mp4; the second one represents a soft behaviour:https://journals.aps.org/prx/supplemental/10.1103/PhysRevX.6.021011/fluid_tissue_v0_0.2_p0_3.8_Dr_0.1.mp4. In both videos, we selected a total of 13 frames with steps of 3.333 seconds (from t = 0 to 40 seconds).

### Graphlets and motifs selection

The different images from the previous section were used to create a graph of cell-to-cell contacts ((Escudero et al., 2011) and **Supplementary Material and methods**) that served as the source for the graphlet analysis (Pržulj, 2007; Pržulj et al., 2004). First, we adapted the graphlet analysis performed by EpiGraph to the nature of our samples (tessellations). Three graphlets were discarded since they were not possible in the context of an epithelial tissue (Fig. 1 and **Fig. S1**). Second, we used the computer program for graphlet identification and calculation ORCA (Orbit Counting Algorithm) (Hočevar and Demšar, 2014), to extract the different conformations of nodes assembling the graphlets, called orbits (Pržulj, 2007). We computed the Graphlet degree Distribution of the 73 given orbits from the 29 graphlets, and then we removed the non-used ones. The reason to remove these graphlets was that they were either redundant or not possible in a planar tissue. On the first case, G5 and G27 were redundant since, in order to achieve G5 in a plane, there must be a centre cell with 4 sides, the same centre cell captured on G27 (Fig 1 and **Fig. S1**). It may occur that more than one cell is inside G5, which could not be captured by G27, still it would be captured by G5 and the chances of encounter this setting would be very low. Regarding the second case, G20, G22 and G25 were not possible to achieve in a planar tessellation since it is assumed the convexity of the cells. Therefore, we removed them.

### Shape index calculation

We have extracted the shape index, as an indicator of rigidity, from each natural and simulated image, based on (Bi et al., 2015). The global shape index in a tissue was measured as the median of the shape index of the individual valid cells. We quantified the cell area and perimeter using Matlab R2014b. We performed the following approach: We captured the vertex coordinates for each valid cell. Then, we calculated the Euclidean distance between each adjacent vertex, and adding all of them, we got the cell perimeter. From these vertices, a polygon was inferred and we calculated its contained area using the “polyarea” Matlab function.

### Statistical analysis

We have estimated the closest CVTn diagram of a given image in terms of the three GDDs measured in EpiGraph (Epi-Hexagons, Epi-Random and Epi-Voronoi5). We computed the centre of the point cloud formed by the 20 randomizations of a particular CVTn diagram as the mean of those twenty images, obtaining a 3D point. Then, we calculated the Euclidean distance between all the CVTn diagrams central points and the three calculated parameters of the input image, obtaining its closest point, which corresponds to its closest diagram. Furthermore, we checked if this image belonged to the closest diagram point cloud using an outlier detection approach. In particular, we tested if the inclusion of the image into a CVTn diagram point cloud would increase or decrease the standard deviation of the original group. We assigned the probability of being an inlier, which is defined as follows:

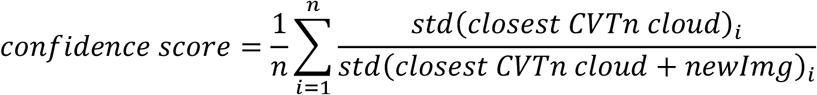

Where n is the total number of coordinates, which in our case is 3 dues to the three-dimensional space; the parameter stands for every different coordinate (Epi-Hexagons, Epi-Random and Epi-Voronoi5); represents the values of the 20 images from the closest CVTn diagram in a specific coordinate and is the value of the input image for the same coordinate. The values range from 0 (very far from point cloud) to +∞ (inside point cloud). We have estimated that with a confidence of > 0.95 the input image is considered to be an inlier.

