## Supplementary Material and methods for "EpiGraph: an open-source platform to quantify epithelial organization"

### Supplementary Information.

#### SUPPLEMENTARY MATERIAL AND METHODS

##### EpiGraph Source code.

The project's code is accessible through Github https://github.com/ComplexOrganizationOfLivingMatter/Epigraph. It is open source and available under GPLv3 license.

##### Pipeline.

The image processing pipeline entails cell recognition, valid cell identification, graph of cell-cell contact creation, polygon distribution analysis and, finally, Epi-Hexagons, Epi-Random and Epi-Voronoi5 calculation. These stages are properly described in the subsequent paragraphs. Moreover, a full set of tutorials explaining how to install and use EpiGraph are available at EpiGraph's wiki (https://imagej.net/EpiGraph). In addition, **Movie S1** summarizes a general example with default options.

- Recognition of existing cells

EpiGraph uses segmented images from natural tissues or simulations as an input. These images must be built in binary format, where one colour should be presented as the background, forming the body of cells, and the other one the cells outline. Each cell is assigned a label using MorphoLibJ (Legland et al., 2016). This identifier allows us to have a record of every cell and its location on the image, which is necessary for the following steps.

- Identification of valid cells

Once we have properly labelled the image, we proceeded to fully analyse it. By default, an invalid region is created, which is initially defined by the boundaries of the image itself. We select all the cells that fall outside of this border and set them as non-valid cells (note that non-valid cells are different from invalid regions, i.e. the latter are not cells). In this way, we avoid incorrect characterization of polygon distribution due to the lack of real neighbours in the margins of the image. However, it is also possible to create a personalized invalid region, by selecting as many cells as you want.

In addition, we define another two kinds of valid cells: the 3-distance valid cells and the 4-distance valid cells. The former represents the cells that do not have a non-valid cell within a distance of three cells connexions. The later kind is formed by the cells having all valid cells in a maximum length of four cells connexions. 7-motifs and 10-motifs require 3-distance valid cells and 17-motifs and 29-motifs 4-distance valid cells.

- Neighbourhood creation

An epithelium can also be considered as a tessellation because there is no space between each cell. Thus, inspecting the number of sides of the cells, we can measure the number of neighbours that will surround it. We, therefore, extend a mask from each pixel of the cell with a given shape and size, both selected by the user. Then, we capture all the cells distinct from the actual cell that fall into this mask and add them as neighbours of the concrete cell. Afterwards, we create a neighbourhood network, modelling each cell as a node and connecting two cells with an edge if they are neighbours.

- Polygon distribution analysis

The polygon distribution is defined by the number of sides of every valid cell. We specify three different areas: the global zone formed by all valid cells, the region in which 3-distance valid cells are contained and the territory in which 4-distance valid cells are placed. For 7-motifs and 10-motifs, we use a path of 4-nodes length (3-distance valid cells as orbits nodes, and any type of valid cell as its branching nodes), and for 17-motifs and 29-motifs the area is defined by the cells contained within the path of 5-nodes length (4-distance valid cells as orbits nodes, and valid cells as its branching nodes). Finally, we calculate the polygon distribution regarding the cells involved in the graphlet calculation.

- Epi-Hexagons, Epi-Random and Epi-Voronoi5 calculation

We have adapted Graphlet degree Distribution agreement Distance (GDD) from Yaveroğlu et al. (Yaveroğlu et al., 2014) to fully integrate it with FIJI java environment. We use GDD to compare two images and describe how similar they are. In particular, minimum distance value (0) means two images are equal, and maximum distance value (1) specifies that they are very different in terms of graphlets. We compute all the graphlets in which the valid cells participate. Then, we use only the graphlets in which 3-distance or 4-distance valid cells are included (depending on the set used). We defined three references to compare with the real images. The first one is the Graphlet degree Distribution agreement Distance Random Voronoi (Epi-Random). A random Voronoi (RV) emerges from 500 seeds randomly placed. Then, we computed the GDD between each one of the 20 RV images and the input image and used the mean of these 20 GDDs as final distance. We also calculated the Graphlet degree Distribution agreement Distance Hexagons (Epi-Hexagons), which measures the difference between a given image and a regular tessellation of hexagons, in a similar way to the Epi-Random. Finally, we computed the Graphlet degree Distribution agreement Distance Voronoi 5 (Epi-Voronoi5) by comparing a given image with 20 diagram 5s from the CVT path and calculating its mean.

- Statistical analysis

Right before adding all the GDD values to the table, we performed a statistical analysis (see **Material and methods, statistical analysis**) to calculate the closest diagram on the CVTn. This test provides a confidence score determining if the source image is part of the CVTn path of reference or, on the contrary, is an outlier.

**Graphical user interface.**

ImageJ's FIJI distribution (Schindelin et al., 2015, 2012) provides a framework, which offers a range of functions and a suitable predefined application programming interface (API). Furthermore, it allows users with no programming expertise to execute and exploit all their functionalities. EpiGraph is prepared for heavy processes. In these cases, some operations such as "calculate graphlets", run in the background, giving the user the possibility to move around the application smoothly. In addition, a progress bar displays the evolution of the task in case it may take longer than expected. FIJI works on the three main Operative Systems (Linux, Mac and Windows) without requiring Java installation since it is usually embedded in the program. FIJI describes different ways it can be operated by external software. One of them is constructing a plugin, which is used by EpiGraph as basic structure to start building. Using Java Swing GUI libraries, we have designed the three windows of the EpiGraph plugin.

<u>- Main Window</u>

The program starts after clicking on *Plugins>EpiGraph*. The first window that appears is the Main Window. It contains a table of processed images (therefore, initially it is empty). Each table row corresponds to a single processing of image and comprises the following columns: Colour, label, Epi-Hexagons, Epi-Random, Epi-Voronoi5, percentage of hexagons, radius of shape, type of shape, kind of graphlet calculation, closest diagram, confidence score and a checkbox. *Colour* column lets you pick the colour of the point from the palette to be visualized later; *Label* shows the name of each image; *Epi-Hexagons*, *Epi-Random* and *Epi-Voronoi5* represent the GDD of the image against hexagons, random Voronoi diagrams and Voronoi 5 diagrams from CVTn, respectively; *Percentage of hexagons* contains the proportion of hexagons for all valid cells involved in graphlet calculation; *Radius* specifies the shape size used to calculate neighbours; *Shape* characterizes the type of form used to calculate neighbourhood; *Kind of calculation* displays which set of graphlets was used; *Closest diagram* represents which number Voronoi diagram from the CVTn is closest to the image; *Confidence score* is the certainty of the image being an inlier of the point cloud of that diagram. Finally, *Select* column lets you choose, by means of a check box, whether you visualise the calculated data in a 3D viewer or not, and in the same way, removes the selected rows if you click on the button *remove rows*. Additionally, you do not need to calculate the graphlet properties every single time. Instead, it is possible to import your own dataset from a properly formatted Excel file, using the *import table* button. Likewise, once you have already analysed several images, you can export your information into an Excel file, using the *export table* button. The exported table includes the polygon distribution of the cells involved in the graphlets calculation. Lastly, you have the option of representing all table rows, by ticking the “true” check box. For that, you should click on the *visualize* button to launch the mentioned 3D viewer.

<u>- Image processing Window</u>

This window will be automatically triggered when a supported file format is correctly selected, after clicking the *open* main window button. This window is composed of a button panel with different processing options, a canvas with the loaded image embedded in it and a polygon distribution legend. The only enabled button when the window is first opened, will be the mode in which you label the image. You can choose to label this image using either 8-connectivity or 4-connectivity and then click on the *label image* button to execute it. After these other buttons become enabled. These buttons are classified in 3 panels:

- *Region of interest*. The main objective of this panel is selecting regions of interest so that only valid regions are processed and invalid ones are discarded. The *create ROI* and *select cells* buttons let you select various rectangular (default) regions and pick individual cells using the FIJI’s ROI manager tool. You can repeatedly combine both options to establish a valid region in which to operate. Furthermore, you can change the ROI selection shape, from rectangular to another shape, using FIJI’s control panel. On the other hand, there is the option to choose an invalid region by clicking over it, after clicking the *add invalid regions* button. This action only lets you store a single set of invalid ROIs at a time, so if you wanted to add an invalid region after saving previous invalid regions, you must delete the former to include a new one.
- *Neighbourhood*. This panel allows you to visualize valid and non-valid cells, invalid regions and polygon distribution values from the current image. To calculate the neighbourhood you should select, using the number selector (located close to *radius*), a size of shape in pixels and a specific geometrical shape using the tool described above (located next to *shape*). This selection depends on the width of the cell outline. If the border cells have a width of 1 pixel, it is enough using a size of 3 pixels to explore the cells vicinity. By default, we expand 3 pixels each cell, using the circular shape, to look for their neighbours. For wider border cells, you can choose higher size of mask with either circular or squared shape. Once

the parameters have been selected, the *test neighbours* button can be pressed to calculate the neighbourhood for the valid cells belonging to the ROI. If the ROI is the whole image, a column next to the legend of polygons shapes, headed by “Graphlets”, would be filled with the polygon distributions of the valid cells. Otherwise, if the ROI is a subsection of the image, a new column appears next to “Graphlets” headed by “ROIs” that will be filled by the polygon distribution of the valid cells belonging to this ROI. Note that in this case, the “Graphlets” column would be filled with the polygon distribution of the all valid cells participating in the ROI graphlets calculation.

Finally, when the columns are filled, an overlay is displayed on the canvas, representing each cell colour coded by polygon number. These colours are the same that the ones at the legend located at the left of the window. In addition, the *toggle overlay* button lets you choose whether to visualize the overlay created by *test neighbours*. The invalid regions and the cells outlines are labelled in black in the overlay, dark grey marks non-valid cells, and the rest of colours (represented in the polygon distribution legend) are reserved for valid cells. It is important to highlight that the bright colours of the legend displayed over the canvas represent the number of sides of the valid cells into the ROI. The same pale colours represent the polygon distribution of valid cells located out of the ROI, yet which still contribute to the graphlet calculation of cells within the ROI. A 4-distance valid cell is a cell that do not have a no valid cell within a distance of four cells connexions. Moreover, a 3-distance valid cells have no valid cells in at least three branched cells.

- *Graphlets*. This panel is designed with the aim of saving graphlet data internally and externally. There is a text box where you can add the data label (image name by default), located after *image label* text. You can also select a colour label for your data, using *pick a colour* button. Finally, you can choose the appropriate method to calculate graphlets data. The options are: 26 graphlets representing 29 cellular motifs (29-motifs), 17 graphlets that are contained in 17 cellular motifs (17-motifs), 9 graphlets on behalf of 10 cellular motifs (10-motifs) and 7 graphlets typifying 7 cellular motifs (7-motifs). 7-motifs and 10-motifs are formed by graphlets of maximum 4 nodes, while 17-motifs and 29-motifs make use of graphlets of maximum 5 nodes. 7-motifs and 10-motifs are most useful when images have a small number of cells, since both require fewer cells than 29-motifs and 17-motifs. Once the method has been selected, you can then click the *calculate graphlets data* button to acquire data for all calculated graphlets. When this process is complete, data are automatically added to the main window table. Furthermore, by clicking on *Export Graphlet data* you can export a ZIP file containing: a JPG image representing the neighbourhood, another JPG image capturing the label of the cells, a CSV file storing all calculated graphlet data and a .sif that represent the neighbourhood network.

The window has a progress bar that estimates the process state. You can modify the image zoom by pressing control and rotating the mouse wheel at same time.

- Visualizing Window

We use a 3d viewer to display our calculated results stored in the main window's table. When a row´s checkbox is ticked, it will be plotted. To develop this window, we have used an open source library named Jzy3D (http://www.jzy3d.org/) that is able to generate different graphical representations. In particular, we make use of the 3d scatter plot class.

This window is displayed after clicking on the *visualize* button (located in the main window), even if your table is still empty. This window is composed of the scatter plot figure located on the left and a set of components to modify the appearance of this figure on the right. The plotted figure is delimited by a 3D box with 3 axes: Percentage of hexagons, Epi-Hexagons and Epi-Random by default. These 3 axes can be replaced by any of the following configurations ($X - Y - Z$ axes) using the drop-down list with the label *Axes of figure* (upper right corner of the window):

1- Epi-Hexagons, Epi-Random, Percentage of hexagons (default)
2- Epi-Hexagons, Epi-Random, Epi-Voronoi5
3- Epi-Hexagons, Epi-Voronoi5, Percentage of hexagons
4- Epi-Random, Epi-Voronoi5, Percentage of hexagons

The Percentage of hexagons axis encapsulate values between 0 – 100, and the others between 0 – 1. These limits can be modified for zooming in on the individual axes. We used three *rangeSliders* to select the range for each axis to be represented due to the limitations of the Jzy3D library. This library only provides you the possibility of zooming the $Z$ axis turning the mouse wheel. You can visualize the three *rangeSliders* (one per axis) just below the *Axes of figure* drop-down list.

By default, the scatter plot displays the CVTn path, shown as individual dots: The darkest dot represents the average of 20 Voronoi diagrams 1 in CVTn and the lightest one is the average of 20 Voronoi diagrams 700 in CVTn. You have the option to disable the visualization of these references clicking on the checkbox 'Show reference' (just below *rangeSliders*), to only display calculated data. One can also adjust the size of the dots by modifying the position of the slider bar, located just below the previously mentioned checkbox. To select the different modes in which graphlet data of the CVTn reference can be calculated, a drop-down list can be deployed by clicking on the label *Motifs of CVTn reference*. This list allows you to represent the CVTn path depending on the method with which the graphlets were calculated: 29-motifs, 17-motifs (by default), 10-motifs and 7-motifs. In this way, you can export the visualized CVTn path by clicking on the button with label "*Export actual CVT reference*" as an excel file with the GDDs to represent yourself your acquired data elsewhere (as in "*Export table*" of the Main Window).

The dots shown in the figure in the Main Window can be modified by changing the colour box in the Main Window's table. In addition, a .png figure screenshot can be saved, by pressing the *Export view* button. For more detail, different angles and modes of visualization are available. By clicking the figure and moving the mouse, one can change the viewing angle, while double clicking on the figure will automatically perform a 360 degrees' rotation.

**Functionalities.**

Along the program execution pipeline, there is the opportunity to develop a set of functionalities:

- Label image

Once you have selected an image to process, this image is binarized and the background is analysed to detect the cells and their outlines. If the number of white pixels is higher than black pixels, white pixels will be considered as cell's body, and vice versa. After that, each cell's body region of the image is assigned with a unique label. Thanks to the extensible architecture of FIJI, in which you can install plugins to add additional functionalities, this process of labelling regions is made using MorphoLibJ functionalities. Specifically, we have used the connected component labelling, which transforms a binary image into a labelled one by assigning a specific number to each connected component.

- Create ROI

Though it is possible to analyse the entire image, you can also process a smaller region of interest (ROI). Through FIJI's Roi Manager we can manage the ROIs, saving and performing operations on one at a time. We have selected two default operations within EpiGraph: rectangle (or any available shape) and multipoint selection. The former creates a rectangle selection (by default) and defines all the cells that fall inside it as valid cells. The latter enables the multipoint function with which it is possible to select individual cells. However, it is also possible to change these default forms of selection, by going straight to FIJI's main window and picking any from the toolbar.

-Select invalid regions

As your image may contain artefacts in the form of false valid cells, we have made it possible to mark these zones as invalid regions. All valid cells surrounding the mentioned invalid region will be considered as "non-valid cells". It is possible to select several areas of the image using the multipoint tool, to convert them to invalid sections. To validate this action, the selected invalid region should have the same colour as the cells' background.

- Testing neighbours

As mentioned the previous section, EpiGraph is able to calculate the polygon distribution using 2 essential parameters: a given pixel radius and an element shape. The element shape expands to a given radius of the cells of interest, looking for cells neighbouring. To ensure that the neighbourhood is correctly captured, we allow the users to verify if the image has the right parameters by themselves. The polygon distribution of the image will appear with numbers at the left side of the window and each cell will also be painted with a colour representing its number of sides. Additionally, non-valid cells are coloured dark grey (almost black). Depending on the method chose to calculate graphlets (29-motifs or 17-motifs considering 4-distance valid cells; 10-motifs or 7-motifs considering 3-distance valid cells), the zone coloured with bright colours will be established only by a group of central cells. In the same way, when we select a ROI, the cells affecting the cells within the ROI will be represented in pale colours. Furthermore, due to the variant number of 3-distance and 4-distance valid cells that are going to be filling the ROI, you may encounter when a series of particularities:

- Some cells of your defined ROI could not be final valid cells (3-distance or 4-distance). If the ROI is not surrounded by a minimum of 4 cells in all directions, the number of 4-distance valid cells is going to be lower than the number of 3-distance valid cells. Thus, if you select 17-motifs or 29-motifs, some cells will not be 4-distance valid cells and will not be used in the final graphlet computation as principal nodes.
- An empty ROI. If you select a region with no 3-distance or 4-distance valid cell, you will obtain an empty neighbourhood.
- The pale coloured cells affecting the 3-distance or 4-distance valid cells will differ, when switching from 17-motifs to 7-motifs (for example). Therefore, a different number of cells will be affecting the bright coloured cells.

Finally, the calculations for neighbourhood will be exploited by *calculate graphlets* module, if parameters are not modified since this step.

- Calculate graphlets

The main function of EpiGraph is the graphlet comparison. It begins by checking if there are any selected cells or ROIs. Then, in case any configuration has changed, we re-compute the neighbourhood, otherwise we take the information from a previous computation. From this neighbourhood and valid cells, it calculates the graphlets for the involved network of neighbours. As mentioned in previous sections, it would be incoherent not to filter the graphlets, so we refine it by adding only the nodes at a fixed distance from the border nodes to the final graphlets. We first calculate the total set of graphlets for valid cells and then select a filter that depends on the chosen type of Graphlets to be implicated: 29-motifs, 17-motifs, 10-motifs and 7-motifs. The involved orbit nodes for graphlet calculations will be referred as 3-distance and 4-distance valid cells. When we have the final graphlets, we calculate the three distances (Epi-Hexagons, Epi-Random and Epi-Voronoi5). Depending on the number of graphlets selected, a variable number of orbits are used in the comparison. Finally, the results will be added automatically to the table on Main Window.

- Statistical analysis

Once all GDD data from an input image have been calculated, a statistical analysis is carried out to check if the GDD values of a certain image matches with the CVTn scale or if the image is out of the CVTn path in terms of organization. First, the closest CVTn diagram to the image is computed, estimating the Euclidean distance by considering 3 dimensions: Epi-Hexagons, Epi-Random and Epi-Voronoi5. Thereafter, it is checked whether the image could belong to the closest diagram CVTn point cloud. At this point, the method generates a confidence score for the comparison to the CVTn (see **Material and methods, statistical analysis**). The closest diagram and the confidence score are computed and added to the main window table immediately upon calculating the GDD values or importing from an existing excel document. Alongside these two parameters, when exporting to an excel, the Euclidean distance to the closest diagram is also presented as a column.

- Visualization

Visualizing results properly is a major feature and an important challenge to interpret results. Thus, we have embedded Jzy3d chart in a Java *JDialog* where you can visualize the calculated results' three coordinates (any combination of Epi-Hexagons, Epi-Random, Epi-Voronoi5 and percentage of hexagons) as a point in a scatter plot. Once the points are represented you can compare them with our CVTn reference. At this point, you may want to change the illustrated CVTn selecting the number of graphlets used on the computations, although it is advisable to compare images with the same configuration (7-motifs, 10-motifs, 17-motifs or 29-motifs). Additionally, it is possible to increase or decrease the size of the dots and zoom each axis manually. Finally, you can export the actual view of the chart to an image file.

**Dependencies.**

FIJI is designed to add functionalities via several routes, one of which is Plugin. In order to have simple control of dependencies and project settings, it uses Maven. Maven is a tool that is broadly used for supervising and building Java-based projects. It integrates several tools, such as Javadoc, to make all the programming steps easy. Dependencies are downloaded and updated when available. Furthermore, it helps you create the package with all your code and zips it in a .jar file that will be the Plugin format of FIJI. All this is achieved through its project object model (POM), which you can shape to your project adding mailing lists, issue tracking and more.

**Supported file formats.**

EpiGraph's input is an image. This image should be properly segmented and grey-scale, however, there are several options within EpiGraph to configure it depending on the type of the image. We allow images with different sizes of borders, since it is possible to increase the radius of the mask, changing the way in which the neighbourhood is built. However, it is mandatory that images are binary images (8 and 32 bits RGB images can be loaded, but they might not be correctly computed), where one colour is presented as the background, forming the body of cells, and the other the cells outline. Regarding the image file extensions, we entrust FIJI with the image opening and the supported files, so individual image file extensions allowed by FIJI would be supported in EpiGraph. We cannot admit any sequence of images and the program is sensitive to single images with high resolution. Accordingly, we warn the user when the images overpass the 3000px either along the height or width, to prevent possible execution problems due to the high computational complexity.

**Quick step-by-step EpiGraph's usage.**

- Installation

*Through FIJI/ImageJ update site:*

The usual way to install a FIJI plugin is through his on-site updater. It's usually located on *"Help > Update FIJI"*. Once it is open, you click on "*Manage updates* *sites*", look for EpiGraph and tick the checkbox next to it. Finally, "*Apply changes*" and you should have successfully installed this plugin. With this option, you can automatically get the latest version of EpiGraph.

*Manually:*

On the other hand, you may just want to download the .jar file from http://bit.ly/EpiGraph-1-0-2 (or even generate your own .jar from source code) and install it manually. To do this, you can click either *Plugins>Install or Plugins>Install PlugIn*. Then you select the provided .jar and you should be able to run EpiGraph.

<u>- Simple example of a complete analysis</u>

Calculate Epi-Hexagons, Epi-Random, Epi-Voronoi5 and percentage of hexagons from a given image and visualize it in the 3D visualizer.

1. Select *open*.
    a. Select a supported image.
    b. If the image is supported, a window with the image will open.
2. Pick connectivity:
    a. Select the connectivity of your image. Usually 8-connectivity.
    b. Press *label.*
3. Default configuration:
    a. Radius of 3 pixels.
    b. Circle shape.
4. Calculate graphlets and statistics:
    a. Write a name for your image.
    b. Pick a colour for your image.
    c. Select *29-motifs* in the combo box.
    d. Press *calculate graphlets*. This will calculate Epi-Hexagons, Epi-Random and Epi-Voronoi5.
    e. When GDDs have been calculated, a statistical analysis is carried out providing a confidence score and the closest CVTn diagram to the image. Once it is finished, if the name box is not empty, all data will be automatically added to the main window table.

5. Export graphlets data:

a. Press the *Export graphlets data* button.
b. EpiGraph create a zip folder containing several files; an image with all the cells labelled with their corresponding identifier, another image representing the polygon distribution and, finally, a '.csv' file with the graphlets for all the valid cells.

6. Visualize your results:
    a. Return to the main window.
    b. Click on *visualize.*
7. Export view:
    a. Staying in the visualizing window, you have the option of exporting the actual view into a '.png' file at every stage.
    b. Press *export view*.

Congratulations! You have done the complete pipeline. Now, check where your image is in regard to the CVTn scale. You should analyse whether or not your image is near the CVTn (reference), which can be done using its closest diagram and confidence score from the table.

- Create region of interest

If you wish to test a sub-region of your images, you can create a region of interest (ROI). We have already opened an image and labelled it within EpiGraph.

1. In the image processing window:
    a. Press *Create ROI*.
    b. The default ROI is square shaped; however, you may want to test another shape. You can now return to the FIJI application and select any of the existing ones on the toolbar.
    c. Click *done*. Your ROI is managed and stored in FIJI Roi Manager.

You have created your ROI; however, you cannot see how it affects the image yet.

2. Press *Testing neighbours:*
    a. An image representing each cell by its number of neighbours will appear as an overlay in your image. Remember that the valid cells in the ROI, which appear with intense colour, are the 3-distance or 4-distance valid cells (depending on your graphlet selection), and the pale colours represent the rest of valid cells that

affect the computation of graphlets. The result will be represented in the polygon distribution legend.

Once this procedure has been accomplished, you can continue the execution in the same way as before: graphlet calculation and statistics, export of results and visualization.

<u>- Combine ROIs</u>

As regions of interest are managed by the ROI manager you can add several ROIs to the image. You have already opened the image and properly labelled it.

1. Press *Create Roi*:
    a. Select a left-hand region with the rectangular ROI shape. You should not select any of the cells on the right-hand side yet.
    b. *Done*.
2. Select individual cells:
    a. Click *Select cells*.
    b. Pick only cells on the right-hand side of the image.
3. Test your setup:
    a. You should be able to see the cells that fall into both ROIs.
    b. Press *test neighbours*.
4. Oh! You realize you have made a mistake:
    a. You do not want the first ROI.
    b. Go to the ROI manager.
    c. Select the first ROI and click Delete.
5. Test the new selection:
    a. You will now represent only the existing ROI on the right.
    b. Click *test neighbours* again.

If you create some ROIs, the selected cells will be all the cells that fall into any of the ROIs. The logical operation would be an OR, so ROIs could be disconnected.

<u>- Selection of the number of motifs</u>

There are 4 possible configurations. In this example we will create an analysis with 29-motifs using 26 graphlets.

1. Open an image representing cells with 4-connectivity and borders with 1-pixel width.
2. Label it:
    a. Click *label.* A warning notice advises that there are very few cells in the image. Your full image is tagged with the same label because all the cells are connected.
    b. Select now 4-connectivity.
    c. Click *label*. That works as expected! Each cell has a different tag.
3. Test neighbours by default.
4. 29-motifs:
    a. This configuration has the maximum number of graphlets present in our study.
    b. Change the combo box to *29-motifs*.
5. Calculate graphlets:
    a. Add a proper name to your image.
    b. Press calculate graphlets to add the full results to the table.
6. Visualizing:
    a. Change the colour of your row to a pink-ish colour. This will let you differentiate your data from the reference, which is black, grey and almost white.
    b. Click on *Visualize*.
    c. As a default, your reference is *17-motifs.* So, change this option in the combo box to *29-motifs.*
    d. Modify the axes of figure to ‘Epi-Hexagons, Epi-Random, Epi-Voronoi5’.
    e. Change the range sliders at your right side to get an adjusted visualization per axis.

You will see that the reference has changed regarding the default 17-motifs reference. Your image is probably aligned to the CVTn reference. This is because your image is within the CVTn path, otherwise the image organization would be different to our model of reference. You can verify this with the confidence score, which also provides the closest diagram to the input image.

- Increase the radius of neighbourhood.

You have a segmented image with a border wide of 3 pixels, and you want to capture the real number of neighbouring of each cell. You have already opened the image and properly labelled it.

1. Test if neighbours are correct:
    a. Since you do not know which pixel radius you should select, test various shapes and radii in order to see if the cells with 6 neighbours are actual 6-sided cells.
    b. Change parameters and click on *test neighbours* to make sure the neighbourhood is correct.

Once your polygon distribution is correct enough, you can follow the next step of the analysis protocol.

- Import/export table

You may want to save your results and to do this you can export your results to an excel file, simply by pressing the button export in the main window.

On the other hand, if you would like to continue your session right where you left off, you could import your exported excel file to the table and carry on with your analysis. You can add as many .xls files to the table as you want and we allow duplicates. If you want to remove them, just tick them at the *Select all* column and click on *Delete rows*.

Another option is to calculate the mean values of a particular set of images outside EpiGraph. To do this, import the excel with the mean values. EpiGraph will automatically calculate all the values of the closest diagram and its associated confidence score, enabling you to improve your analysis on that dataset.

Troubleshooting

If the visualization is not working properly, you may need to update FIJI after the installation.

If you find something is not working with any functionality of the application, you can send an e-mail to. Reporting bugs can also be done through Github https://github.com/ComplexOrganizationOfLivingMatter/Epigraph/issues. Known issues are held at the same page.

**License Information.**

To encourage the sharing of resources, EpiGraph is published under an open-source (GPLv3) license, which can be downloaded from https://github.com/ComplexOrganizationOfLivingMatter/Epigraph/blob/master/LICENSE.

**SUPPLEMENTARY FIGURES**

**Figure S1. Graphlets and orbits configuration.** Illustration of graphlets networks used in (Pržulj, 2007). Each graphlet configuration is labelled with $G_n$, in which, 'n' is the graphlets number (from $G_0$ to $G_{29}$). These labels match with the cellular motifs in (**Fig. 1**). Each vertex represents a cell and each edge the connection between two cells. Therefore, the graphs represent the connectivity network for each matched cellular motif. The four graphlets labelled in red ($G_{20}$, $G_{22}$, $G_{25}$ and $G_{27}$) are the ones discarded in our work. The digits (from 0 to 72) at some mauve vertices indicate the orbits number counted for the GDD calculation.

**Figure S2. Pipeline for the Graphlet degree Distribution agreement Distance (GDD) calculation.** Scheme representing the protocol sequence to calculate the GDD between two segmented images: Eye and Diagram 1. First, a network of cell-to-cell contacts is computed defining the centroids of valid cells as nodes and its connections with neighbouring cells as edges. The nodes are represented with the same colour code as in **Fig. 2**. Second, the graphlets are extracted from the network. This enables the calculation of an index of the distribution of every graphlet. Finally, a comparison between the two Graphlet degree Distributions is performed to obtain the GDD value.

**Figure S3. CVTn scale capturing tissue organization.** Representation showing how the CVTn iteratively evolves (**Material and Methods**) from a disordered and heterogeneous tessellation (V1) to a more homogeneous and uniform Voronoi (V700). Each diagram is presented in circles surrounded with greyscale tones. The series of dots between the circles represent the middle diagrams amid V4 and V700. CVTn, based on the original CVT path**,** has been shown to be a good descriptor for identifying a tissue in homeostatic state. The graphic displays the resulting relationship between the natural samples and their most akin Voronoi diagram (signed by a double arrow) in terms of organization computed by EpiGraph. In addition, it shows which biological samples were contained within the scale (IN) and which were not (OUT). The colour code of the actual samples is to the same as that in **Fig. 2D.**

**Figure S4. Comparisons between epithelia and CVTn path.** 5-dimensional scatter plot representing every possible comparison between Epi-Hexagons, Epi-Random, Epi-Voronoi5, percentage of hexagons and shape index. The X and Y axes for each chart correspond to its column and row names respectively. The CVTn path is shown starting at diagram 1 until diagram 100 (from 1 to 20, from 30 to 100 by steps of 10), and they are represented in greyscale beginning in black and reducing its darkness with the increase of the diagrams. A set of natural epithelia are display: cNT (16 samples, light blue), dWL (15 samples, green), dWP (16 samples, red), Eye (3 samples, orange) and dMWP (3 samples, violet). The mean value is represented as a circle and their individual values as smaller circumferences. In the shape index column, values up to 3.81 represent solid tissues, painted in orange; shape index values greater than 3.81 indicate fluid tissues, in green. The histograms represent the dots density along each column of comparisons. These graphs complement **Fig. 2**.

**Figure S5. Comparisons of the different simulations regarding the CVTn and natural tissues.** All the possible combinations from Epi-Hexagons, Epi-Random, Epi-Voronoi5, percentage of hexagons and shape index in 2-dimensional graphics. The CVTn path is shown starting at the diagram 1 until the diagram 100 (from 1 to 20, from 30 to 100 by steps of 10). The average values of 20 replications per diagram, are represented as greyscale dots beginning in black and reducing its darkness with the increase of the diagrams. Images from biological samples are marked as dots representing their mean: cNT (16 samples, light blue), dWL (15 samples, green), dWP (16 samples, red), Eye (3 samples, orange) and dMWP (3 samples, violet); Simulations are represented with their mean (circle) and their individual values (circumference): Proliferative Control (20 replicates, carnation pink), Case III (17 replicates, hot pink) and Case IV (15 replicates, purple); Non-proliferative control (20 replicates, blue bell), Squared (20 replicates, azure blue) and Elongated simulations (20 replicates, Cornflower Blue). In the shape index column, values up to 3.81 represent solid tissues, painted in orange; shape index values greater than 3.81 stand for fluid tissues, in green. The histograms represent the dots density along each column of comparisons. These graphs complement **Fig. 3**.

**Figure S6. Shape index values for the analysed images.** Plot with shape index and Epi-Random as axes. The orange region represents the shape index values that define a tissue as ‘solid’, and the green zone as ‘fluid’. Circles are the average value obtained from the individual samples from the natural images: Eye, cNT, dWL, dWP and dMWP; CVTn (from diagram 1 to 100), Proliferative Control, Case III and Case IV; Non-proliferative control, Squared and Elongated simulations. Triangles stand for the average value from solid/fluid tissues: WT, (blue) and *Mbs-RNAi* (orange).

#### LEGENDS OF SUPPLEMENTARY FILES

**Movie S1. A general example of the usage of EpiGraph.** The movie shows all the functionalities of Epigraph and how to use them, in general terms.

**Table S1. GDD values between pairs of images from images of biological samples and Voronoi Diagrams.** Graphlet degree Distribution agreement Distance mean between each row and column is shown. Data is divided by the used cellular motifs (17-motifs and 29-motifs). The used samples are Voronoi 1, 4 and 5: 20 replicates; dWP: 16 replicates; dWL: 15 replicates; Eye: 3 replicates.

**Table S2. GDDs and percentage of hexagons of CVTn reference using all the cellular motifs sets.** Data are distributed depending on the cellular motifs used (7-motifs, 10-motifs, 17-motifs, 29-motifs). Mean and standard deviation of percentage of hexagons, Epi-Hexagons, Epi-Random and Epi-Voronoi5 are shown, along with their associated diagram. 20 replicates of each diagram are represented.

**Table S3. Outlier detection results of natural images, simulations and rigid/soft tissues.** For the mean of each set of images, the closest diagram and confidence score are calculated (see Material and methods, statistical analysis). In green, the confidence score above 0.95 corresponding to inliers. In red are marked the confidence scores below 0.95, which corresponds to outliers.

**Table S4. Measurements of fluidity from simulations and actual epithelia using the shape index.** The shape index of all the tessellations used along the manuscript have been calculated in terms of the average of its median and mean values for each sample. The green colour is referred to a tissue with a shape index defined as fluid, and the orange colour is identified as solid.
