## Supplementary figures and images for "EpiGraph: an open-source platform to quantify epithelial organization"

### Fig. S1

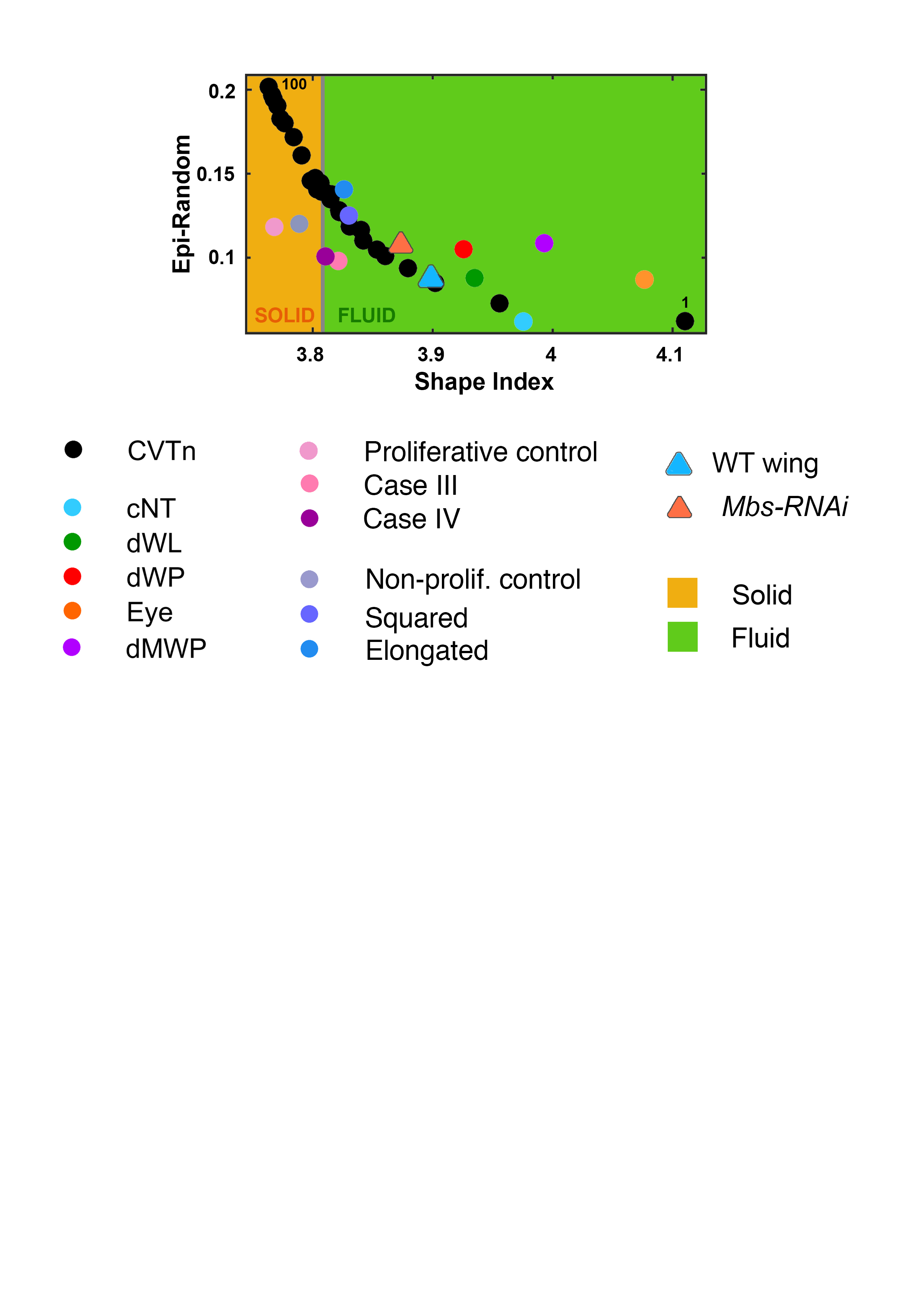

### Movie S1

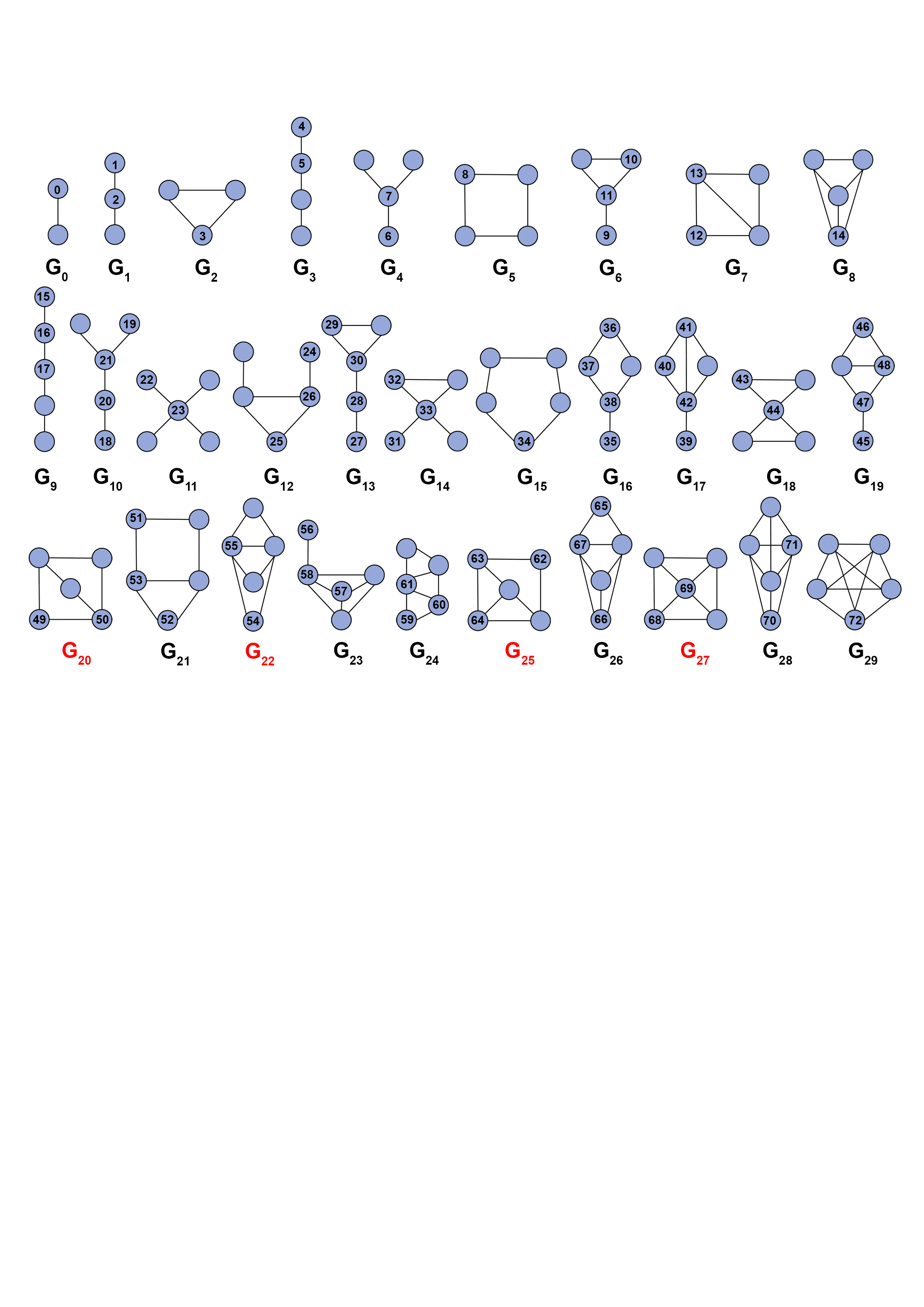

### Table S1

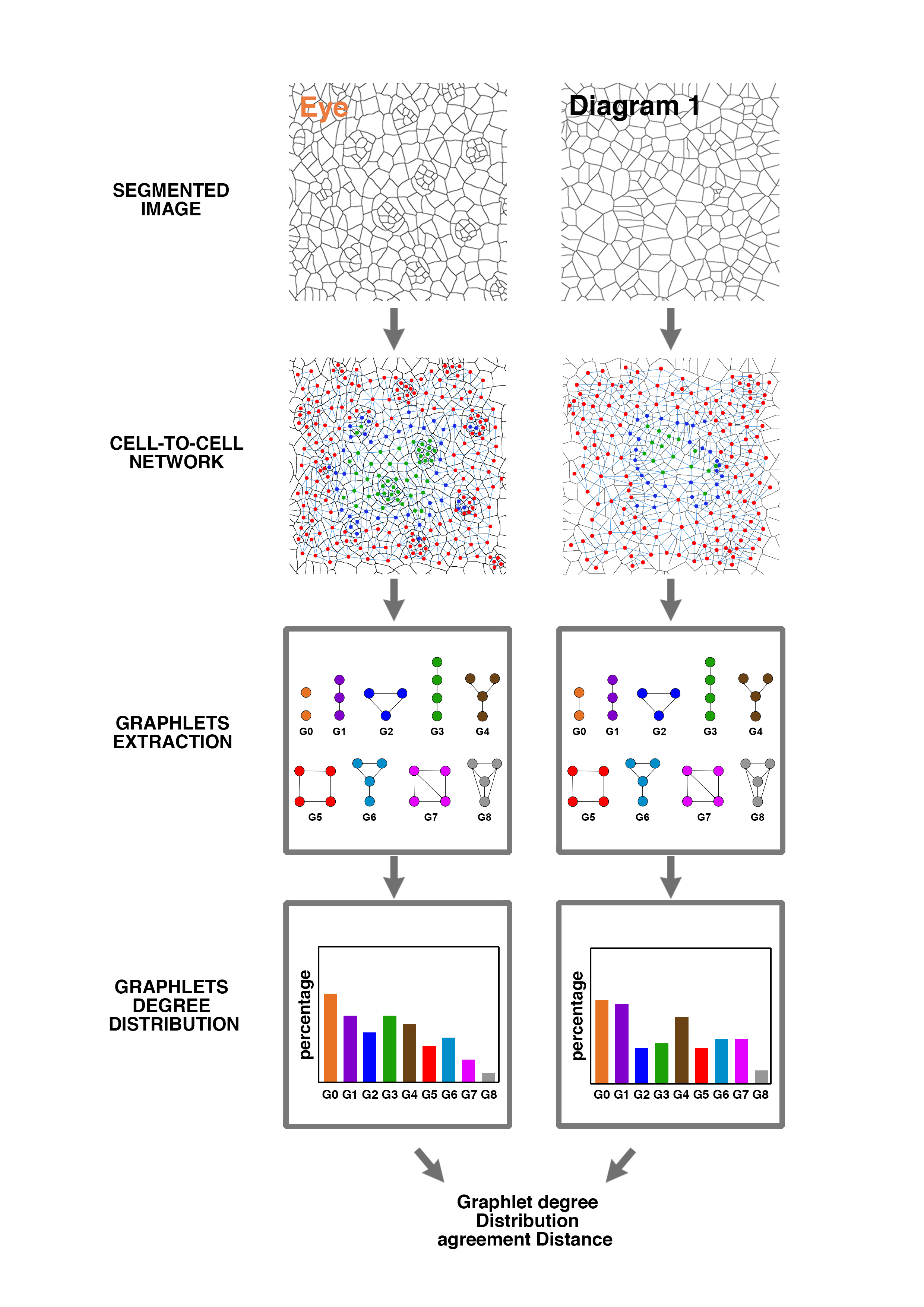

### Table S2

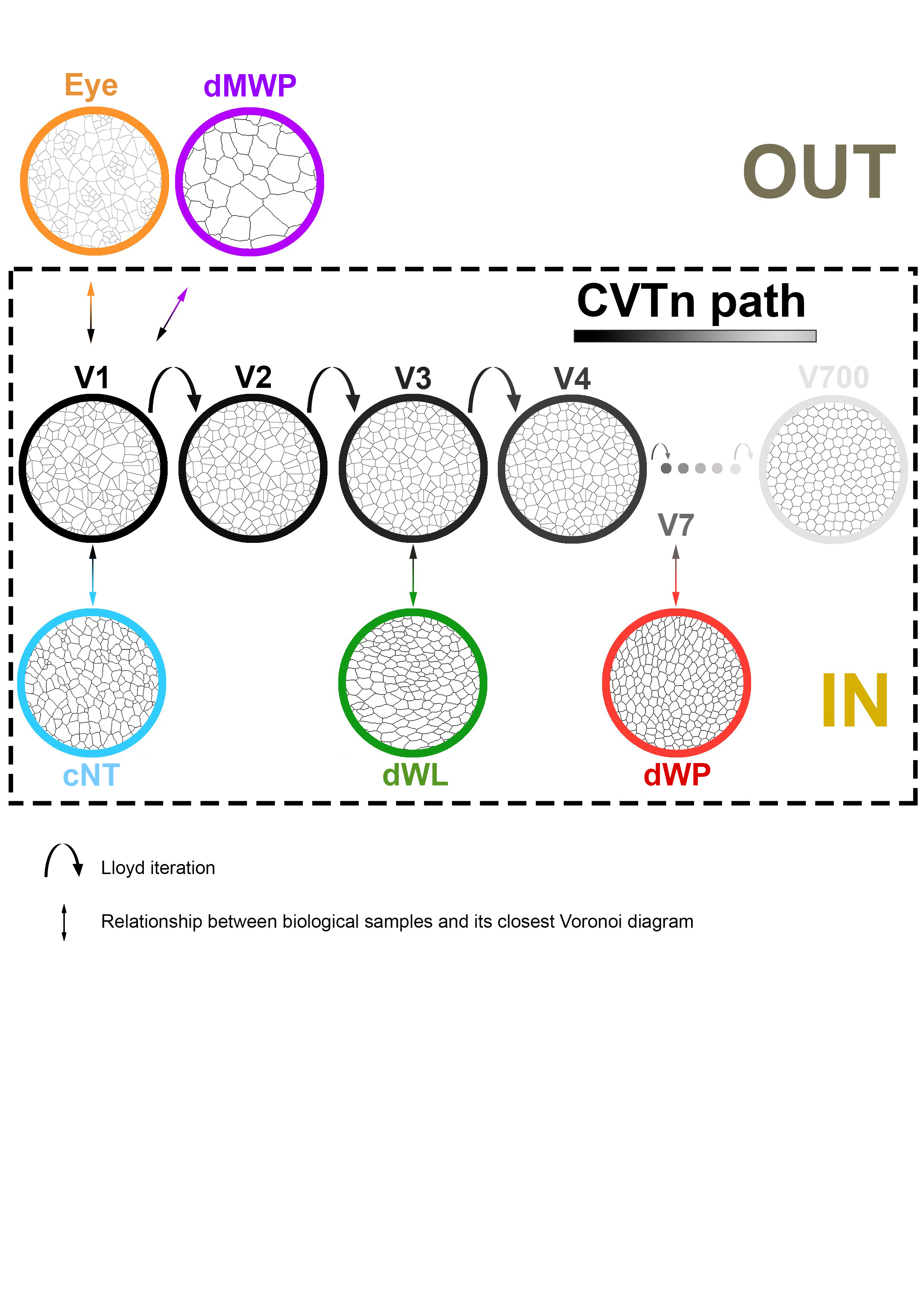

### Table S3

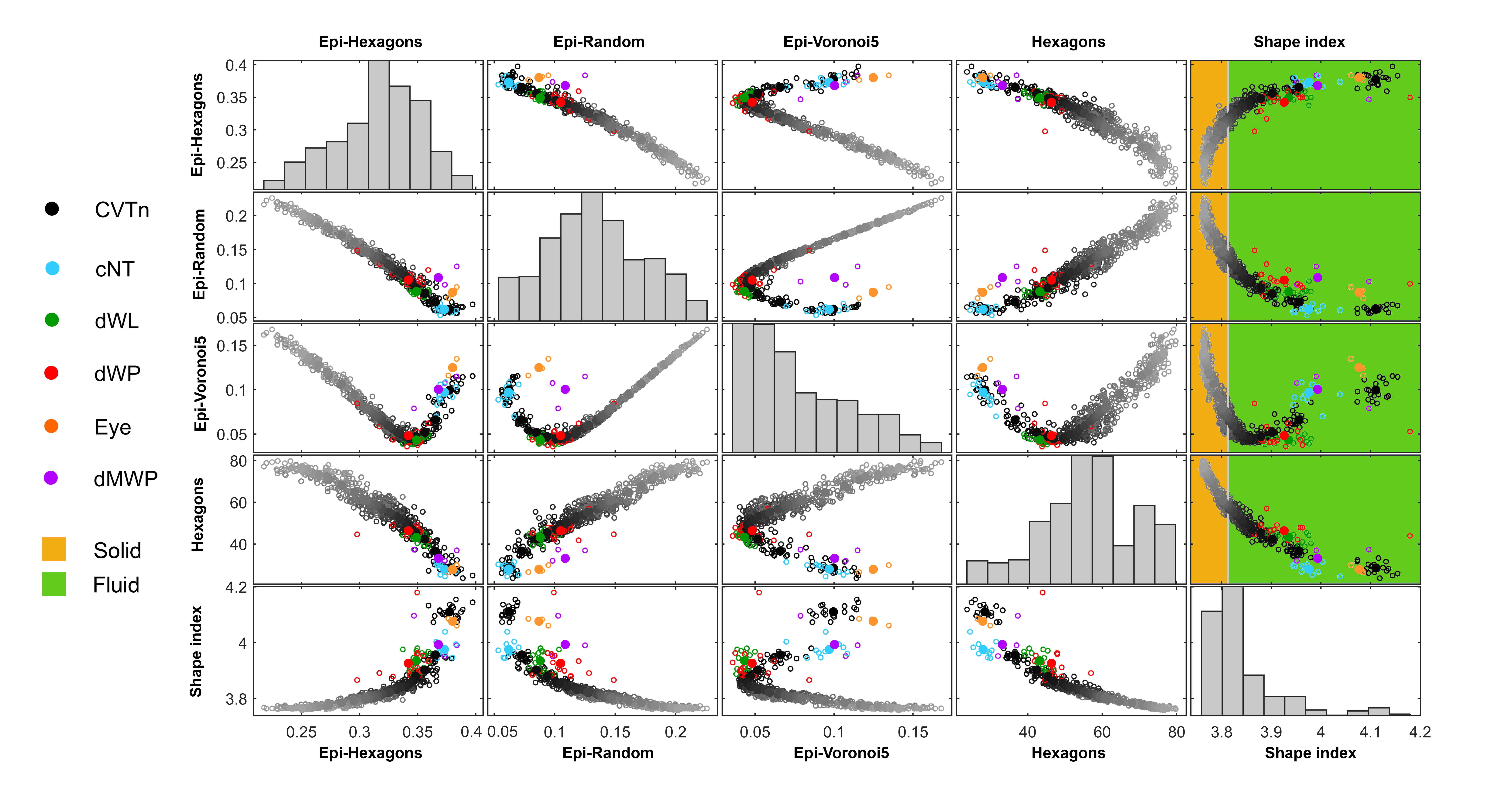

### Table S4

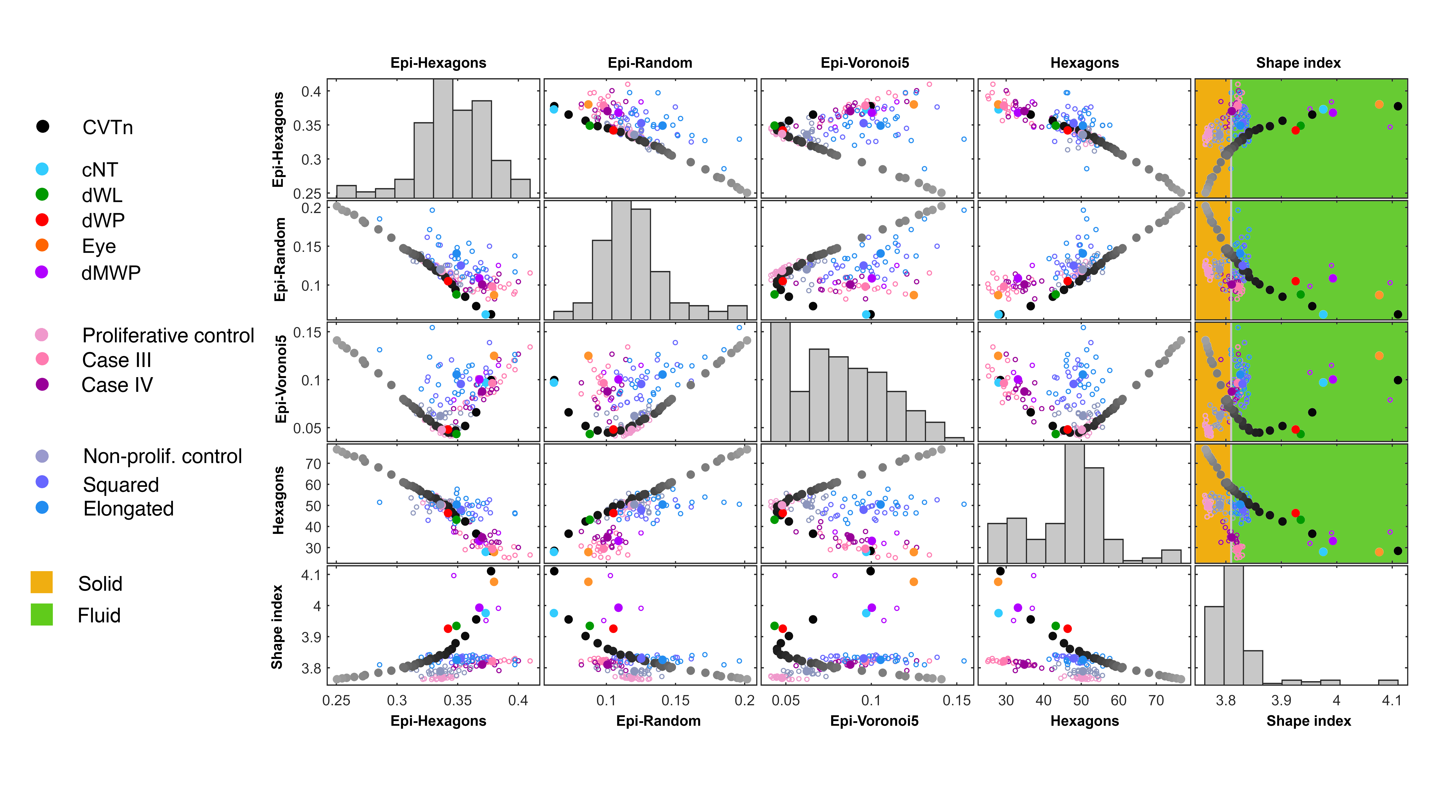
